# Broad-spectrum antiviral inhibitors targeting pandemic potential RNA viruses

**DOI:** 10.1101/2023.01.19.524824

**Authors:** Gustavo Garcia, Joseph Ignatius Irudayam, Arjit Vijay Jeyachandran, Swati Dubey, Christina Chang, Sebastian Castillo Cario, Nate Price, Sathya Arumugam, Angelica L. Marquez, Aayushi Shah, Amir Fanaei, Nikhil Chakravarty, Shantanu Joshi, Sanjeev Sinha, Samuel W. French, Mark Parcells, Arunachalam Ramaiah, Vaithilingaraja Arumugaswami

## Abstract

RNA viruses continue to remain a clear and present threat for potential pandemics due to their rapid evolution. To mitigate their impact, we urgently require antiviral agents that can inhibit multiple families of disease-causing viruses, such as arthropod-borne and respiratory pathogens. Potentiating host antiviral pathways can prevent or limit viral infections before escalating into a major outbreak. Therefore, it is critical to identify broad-spectrum antiviral agents. We have tested a small library of innate immune agonists targeting pathogen recognition receptors, including TLRs, STING, NOD, Dectin and cytosolic DNA or RNA sensors. We observed that TLR3, STING, TLR8 and Dectin-1 ligands inhibited arboviruses, Chikungunya virus (CHIKV), West Nile virus (WNV) and Zika virus, to varying degrees. Cyclic dinucleotide (CDN) STING agonists, such as cAIMP, diABZI, and 2’,3’-cGAMP, and Dectin-1 agonist scleroglucan, demonstrated the most potent, broad-spectrum antiviral function. Comparative transcriptome analysis revealed that CHIKV-infected cells had larger number of differentially expressed genes than of WNV and ZIKV. Furthermore, gene expression analysis showed that cAIMP treatment rescued cells from CHIKV-induced dysregulation of cell repair, immune, and metabolic pathways. In addition, cAIMP provided protection against CHIKV in a CHIKV-arthritis mouse model. Cardioprotective effects of synthetic STING ligands against CHIKV, WNV, SARS-CoV-2 and enterovirus D68 (EV-D68) infections were demonstrated using human cardiomyocytes. Interestingly, the direct-acting antiviral drug remdesivir, a nucleoside analogue, was not effective against CHIKV and WNV, but exhibited potent antiviral effects against SARS-CoV-2, RSV (respiratory syncytial virus), and EV-D68. Our study identifies broad-spectrum antivirals effective against multiple families of pandemic potential RNA viruses, which can be rapidly deployed to prevent or mitigate future pandemics.

## INTRODUCTION

The ongoing severe acute respiratory syndrome coronavirus 2 (SARS-CoV-2) pandemic exposed the limitations and vulnerabilities of humanity from a lack of preparation to rapidly respond to a large-scale outbreak. It is difficult to accurately predict the causative agent of the next pandemic. However, based on recent epidemics over the past two decades, global climate change, the millions of individuals affected, and the continuously-evolving nature of the RNA genome, arthropod-borne viruses are priority candidates. Vector-borne RNA viruses belonging to the families *Togaviridae –* Chikungunya virus (CHIKV), and *Flaviviridae*, such as Dengue virus (DENV), West Nile virus (WNV) and Zika virus (ZIKV), have had track records of causing epidemics^1-3^. Daytime feeding mosquitoes (*Aedes aegypti* and *Aedes albopictus*) are responsible for the transmission of these viral agents. The virus enters the body through a mosquito bite on the skin, inoculating the host. The virus then replicates within skin cells, specifically fibroblasts and dendritic cells (DCs). After replicating locally, CHIKV disseminates throughout the body, affecting many regions, including the lymph nodes (stromal cells, DCs and macrophages), skeletal muscle (satellite cells and fibroblasts)^4,5^, joints (synovial fibroblasts), brain, and liver. As a result, CHIKV infection can cause skin lesions, severe joint and muscle pain, and headaches, among other symptoms. However, due to similarities to other mosquito-borne illnesses, Chikungunya disease is often misdiagnosed as Dengue fever or Zika disease. Cardiovascular issues may also manifest from CHIKV infection, most often seen in the form of myocarditis^6^. While symptoms can be serious, CHIKV infection is rarely lethal. ZIKV infection in women during pregnancy not only causes preterm birth and miscarriage, but also causes infants to be born with microcephaly and congenital Zika syndrome^7,8^. WNV, another member of the *Flaviviridae* family, is the most common arbovirus found in the United States, causing the development of serious, sometimes fatal, illnesses affecting the central nervous system, leading to encephalitis or meningitis^9,10^.

Given their already-demonstrated epidemic potential, finding effective broad-spectrum treatments against these viruses is of the utmost importance as they become potential agents for pandemics. Mutations in the CHIKV Envelope gene (e.g., E1-A226V, E1-K211E, E2-V264A, E1-I317V)^11,12^, particularly in the Indian Ocean Lineage of the virus, have significantly increased CHIKV adaptability and infectivity. Understanding CHIKV is particularly important because it has already shown the potential to spread like wildfire in the event of even one of these mutations, particularly the E1-A226V adaptive mutation which improved viral replication and transmission efficiency. We have seen the impact of these CHIKV mutations in epidemics and mass outbreaks in Eastern Africa^12^, South^13^ and Southeast Asia^14^, and South America^15^. South America, having served as a hotbed for the spread of a new variant of ZIKV in 2016^16^, knows the impact that these viruses can have in terms of rapid spread and overwhelming medical infrastructure and public health efforts. ZIKV introduced the novel severe disease presentation of fetal microcephaly in infected mothers^16^, the neurological consequences of which are a growing concern seven years later as children born during that time now enter the population as schoolchildren. With the ever-present threat of climate change, the permissible habitat of mosquito vectors (*Aedes albopictus, Aedes aegypti*, and *Culex* sp.)^17^, has expanded, increasing the population that could be readily exposed to the virus^18^. We have seen this many countries, including the United States, where WNV has become endemic due to increasing winter temperatures, precipitation, and drought^18^.

Currently, there are no effective vaccines or therapies approved against these important pandemic potential pathogens. Thus, it is critical now more than ever to find a broad spectrum of antiviral molecules that can be deployed either prophylactically or therapeutically in the event of an outbreak or public health emergency to target the host-cell machinery and stop critical steps of viral infection and replication. The mammalian innate immune pathway, more specifically the type I interferon (IFN) response, is the critical first line of defense against virus entry and replication. Pattern-recognition receptors (PRRs) recognize pathogen-associated molecular patterns (PAMPs) to induce immune responses against invading pathogens. There are four main classes of PRRs that can be split into two groups: 1) membrane-bound PRRs and 2) cytosolic PRRs. Toll-like receptors (TLRs) and C-type lectin receptors (CLRs) comprise the membrane-bound PRRs^19-23^. TLR3, TLR7, TLR8, and TLR9 recognize hallmark genomic components of viruses, more specifically dsRNA, ssRNA, and CpG DNA^19-21^. CLRs can either stimulate the release of proinflammatory cytokines or inhibit TLR-mediated immune action^22^. CLRs are made up of a transmembrane domain containing a carbohydrate-binding domain^19^, which can recognize surface carbohydrates on viruses, bacteria, and fungi^19^. Dectin-1 and Dectin-2 are common immunoreceptor tyrosine-based activation motif (ITAM)-coupled CLRs, which recognize β-glucans from fungi^23^.

Within cells, the RIG-I-like receptor (RLR) and NOD-like receptor (NLR) families make up the cytosolic PRRs. RLRs are composed of three proteins: RIG-I, melanoma differentiation-associated gene 5 (MDA5), and DExH-Box Helicase 58 (DHX58/LGP2)^24,25^. RIG-I is an important mediator of antiviral response against CHIKV and DENV^26^. As opposed to TLRs, RLRs are localized in the cytoplasm. Like some TLRs, RLRs recognize dsRNA produced both as a primary genomic component of viruses and as a replication intermediate for ssRNA viruses^19^. NLRs are a primary component of the inflammasome, a protein complex that activates caspase-1 and the processing of pro-inflammatory cytokines IL-1β and IL-18^27^. NOD1 and NOD2, which have caspase activation and recruitment domain (CARD) motifs, activate NF-κB upon recognizing bacterial peptidoglycans, diaminopimelic acid, and muramyl dipeptide, respectively^19,28^.

The cGAS-STING pathway is an essential component of the innate immune system that detects the presence of non-nucleosomal viral, bacterial or damaged cellular cytosolic DNA. The protein cGAS (cyclic GMP-AMP Synthase) is a signaling enzyme that directly binds cytosolic DNA and dimerizes upon binding to form a cGAS-DNA complex that catalyzes the synthesis of 2’,3’-cGAMP from ATP and GTP^29,30^. The STING (STimulator of INterferon Genes) protein dimer, a membrane receptor on the endoplasmic reticulum, binds this newly formed 2’,3’-cGAMP and then recruits TANK-binding kinase (TBK1) which, in turn, phosphorylates interferon regulatory factor 3 (IRF3). Several other dsDNA sensors also recognize foreign DNA (DDX41, IFI16) and this signals through interactions with cGAS to activate STING. Simultaneously, STING activation leads to IKK activation and the consequent phosphorylation and degradation of IκBα. This results in NF-κB1 relocating to the nucleus, along with phospho-IRF3. Subsequently, type I IFN genes and inflammatory genes are expressed, leading to antiviral and inflammatory responses. In this study, we tested the antiviral activity of various innate immune agonists that target PRRs such as TLRs, STING, NOD, Dectin, and cytosolic DNA or RNA sensors. After an initial screen in human fibroblasts cells, we validated our strongest hits that could inhibit multiple families of viruses in human cardiomyocytes and mouse model.

## RESULTS

### Primary Screen to Identify Broad-Spectrum Antivirals

Utilizing natural and synthetic agonists targeting various PRRs, we performed a primary drug screening (Figure 1) to identify broadly-acting compounds targeting members of *Togaviridae* (CHIKV) and *Flaviviridae* (ZIKV and WNV). A small library of 27 agonists were used for the primary screening. Human fibroblast cells (HFF-1) were treated with four doses of antiviral compounds in triplicate 24 hours prior to infection with the above indicated viruses (Figure 1B). In parallel, drug-treated cells were subjected to viability assays to determine drug toxicity in order to select and determine non-toxic effective doses (Supplementary Table 1 and Supplementary Figure 1). We included IFN-β as a positive control. At 48 hours post-infection (hpi), infected cells were examined for viral-mediated cytopathic effect (CPE). We observed that the following 8 compounds prevented CPE: diABZI, scleroglucan, 2’,3’-cGAMP, 3’,3’-cGAMP, cAIMP, Poly(I:C) HMW, Poly(I:C) LMW, and TL8-506 (Supplementary Figure 1B). At 48 hpi, infected cells were fixed and immunostained with virus-specific envelope antibodies: anti-E1 protein of CHIKV (mAb 11E7) and anti-flavivirus group E protein (mAb D1-4G2-4-15). Immunohistochemistry (IHC) images obtained from each well were then quantified and percent infectivity of individual compound-treated cells was obtained (Figure 1C). We found that the cells treated with the following cyclic dinucleotide (CDN) STING agonists showed below-detectable infectivity at the highest non-toxic dose (100 µg/ml) tested against CHIKV, WNV and ZIKV: 2’,3’-cGAMP, 3’,3’-cGAMP, and cAIMP (Figure 1C). Compounds such as scleroglucan (Dectin-1 agonist), c-di-AMP (bacterial CDN STING agonist), imiquimod (TLR7 agonist) and LTA-BS (Gram-positive bacterial cell wall component) showed moderate antiviral activity (>50% inhibition) against all three viruses. Compounds including c-di-GMP (bacterial CDN STING agonist), MDP (NOD2 agonist), Poly(I:C) LMW, and imiquimod (TLR7 agonist) exhibited potent inhibition of WNV and ZIKV, but not against CHIKV. In some instances, we observed compounds that enhanced viral replication, including LPS-RS, adilipoline, FLA-BS, and c-di-GMP (Figure 1C). Interestingly, although cells treated with the synthetic liposaccharide Pam3CSK4 were healthy across all tested concentrations, we found a dose-dependent enhancement of cell death in CHIKV-infected cells (Supplementary Figure 2A-B).

**Figure 1.**
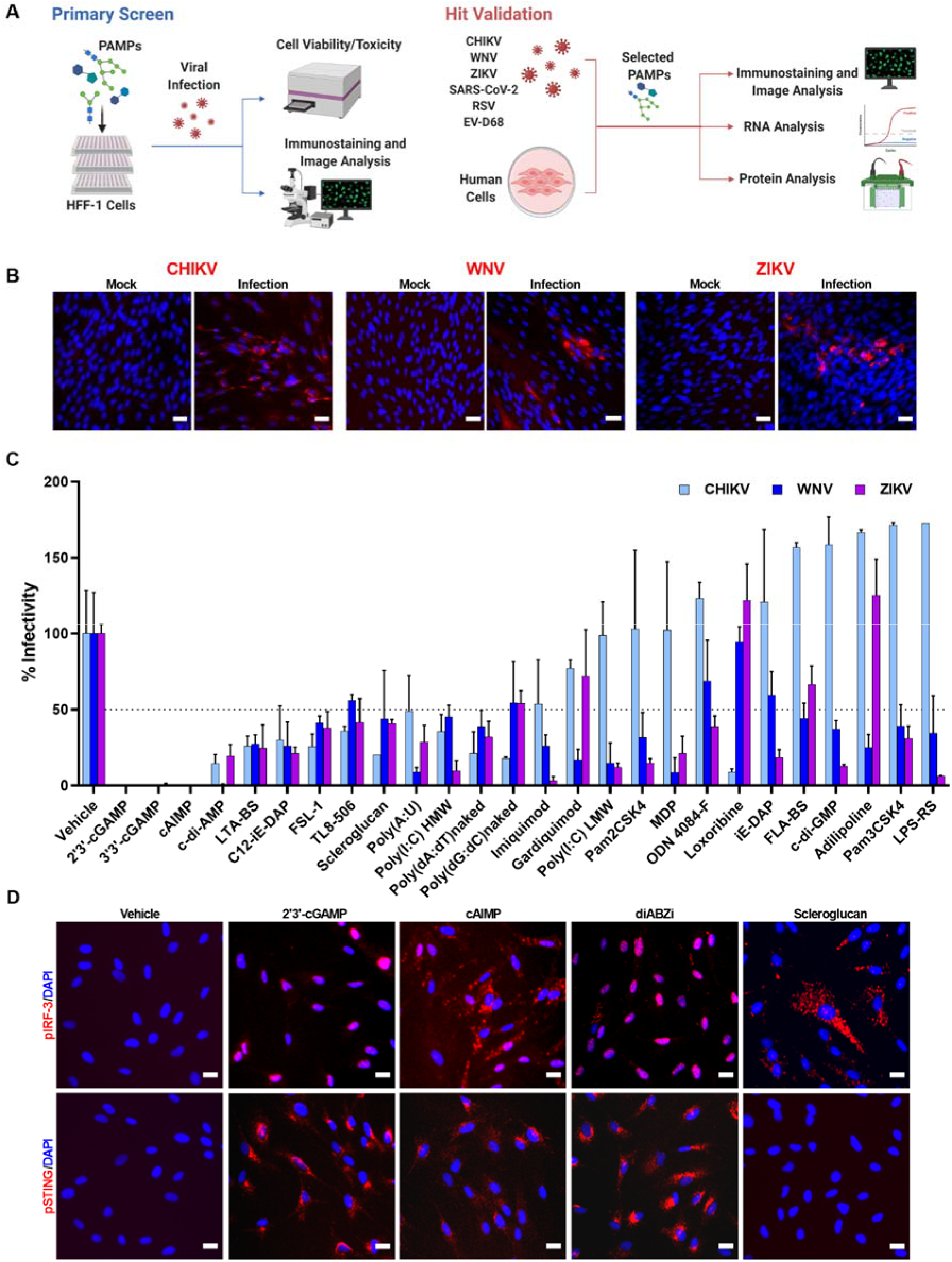
Broad spectrum antiviral screen. (A) Schematic of antiviral screen performed using molecules modulating PAMPs. (B) Immunofluorescence images of human fibroblasts (HFF-1) infected with indicated viruses are shown. CHIKV (MOI 0.1) and ZIKV (MOI 0.1) or WNV (MOI 1) were detected by anti-E1 protein and anti-Flavivirus E protein antibodies, respectively. Scale bar=25 µm. (C) Graph shows the percent infectivity of various compounds targeting CHIKV and flaviviruses. Horizontal dotted line indicates 50% infectivity. (D) Immunofluorescence images show phospho-IRF3 (S386) and phospho-STING (S366) in fibroblast after 2-hour post-stimulation with indicated compounds. Note: Scleroglucan stimulated cells do not have detectable level of phospho-STING, and only few cells have cytoplasmic form of phospho-IRF3. Scale bar=10 µm.

Based on the primary screen hits, we subsequently examined the biological activities of STING agonists in activating the STING-IRF3 and STING-NF-κB pathways, observing that STING agonists specifically induced phosphorylation of STING (Ser366) and IRF3 (Ser386) in HFF-1 cells after 2 hours post-stimulation (Figure 1D), whereas scleroglucan-stimulated cells did not have detectable levels of phospho-STING. All these compounds increased NF-κB p65 form in treated cells (Supplementary Figure 1C). Taken together, we conclude that the STING agonists exhibited a broad spectrum of antiviral activity against RNA viruses by stimulating innate immune pathway.

### Validation of Antiviral Hits from the Primary Screen

Subsequently, we carried out a secondary confirmation using a sensitive RT-qPCR assay to evaluate genome replication of these arboviruses in drug treated cells (Figure 2A). Our data demonstrated that the selected STING agonists (diABZI, cAIMP, and 2’3’-cGAMP), TLR7 agonist (imiquimod), and scleroglucan significantly reduced viral genome replication, which was also confirmed by quantifying viral titer (Supplementary Figure 2). Thereafter, a dose-response study was used to determine the half-maximal inhibitory concentration (IC_50_) of antiviral compounds in human fibroblasts (Figure 2B). We focused on CHIKV, a less well-characterized RNA virus, for subsequent validation of PRR agonists, particularly ligands for STING, TLR3, and Dectin-1. STING is the downstream signaling activator of cGAS-mediated cytosolic DNA sensing pathway and TLR3 (dsRNA) is an endosomal RNA sensor^31^. Compounds 2’,3’-cGAMP, Poly (I:C) HMW, scleroglucan and cAIMP demonstrated antiviral activity at an IC_50_ <500 ng/ml (Figure 2B). Positive control IFN-β and drug-like non-cyclic dinucleotide STING agonist diABZI showed an IC_50_ of 19.69 ng/ml and 10.06 nM, respectively. We also observed that post-treatment of CHIKV-infected cells (2 hpi) with cAIMP and diABZI demonstrated potent antiviral activity, as well (Supplementary Figure 2). Consistent with virulent CHIKV results, a CHIKV vaccine strain (181/25) was also inhibited by STING agonists in the HFF-1 cells (Supplementary Figure 2). Representative IHC images of drug-treated and CHIKV-infected fibroblasts at an effective antiviral dose are shown in Figure 2C. To understand the mode of action of these antiviral compounds, we performed a comprehensive analysis of the innate immune signaling pathway using western blots of HFF-1 cells stimulated by selected STING agonists either with or without CHIKV infection (MOI 0.1) at 24 hpi (Figure 2D). In the uninfected cells, we found that pretreatment with the STING agonists cAIMP, diABZI, 2’,3’-cGAMP, and 3’,3’-cGAMP induced phosphorylation of IRF3. Notably, the cells pretreated with STING agonists had a reduction in total IRF3 levels at 24 hpi. IFN-β strongly induced phospho-TBK1 (Ser172) and phospho-STAT1 (Tyr701). Compared to the STING agonists, scleroglucan-treated cells had only the basal level of phospho-IRF3, indicating specificity and differential mode of actions between agonists and their associated PRRs. CHIKV nsP3 protein level was consistent with antiviral activity of the tested compounds. Taken together, we conclude that STING-mediated activation of the TBK-IRF3 signaling cascade exerts potent antiviral activity against CHIKV.

**Figure 2.**
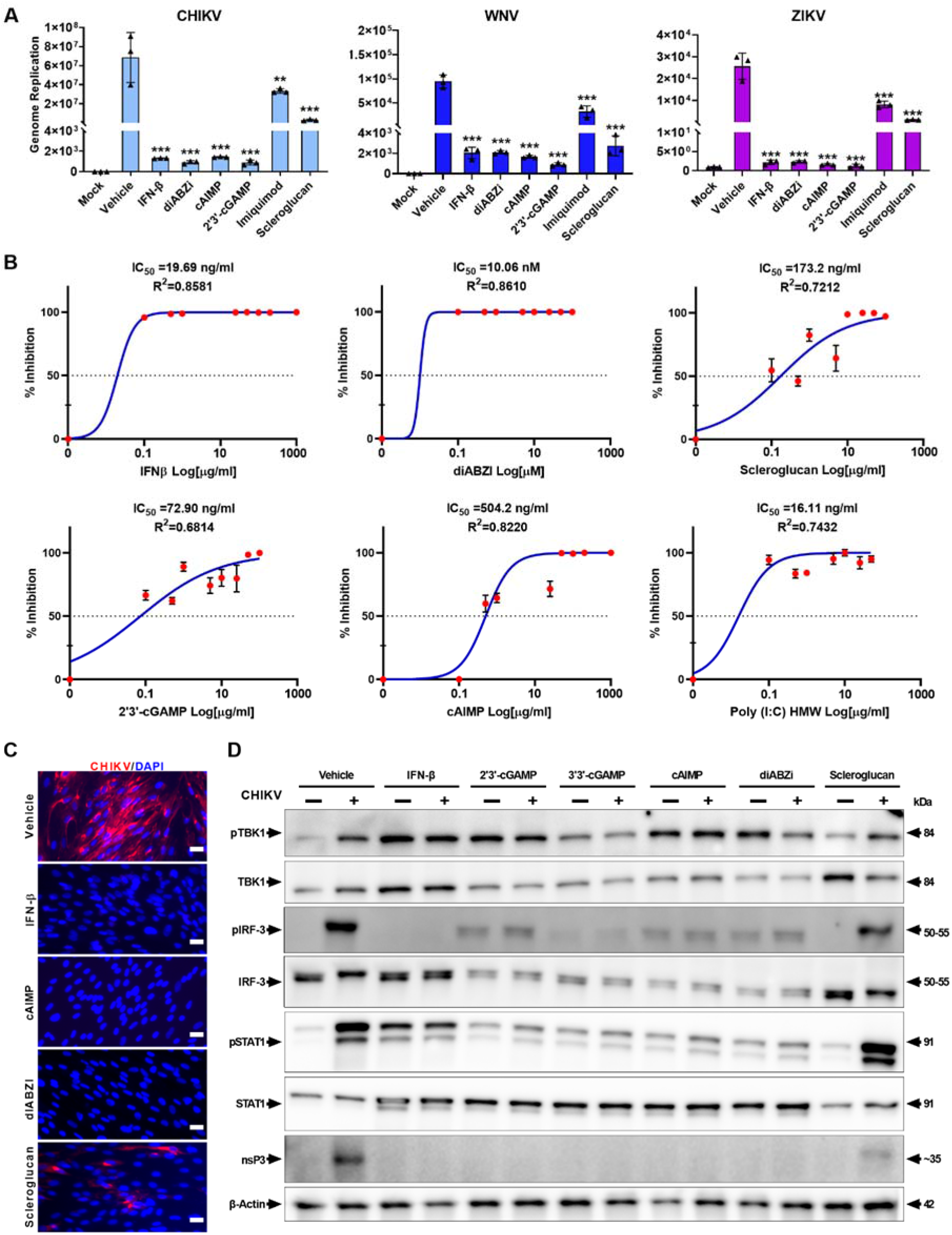
Hit validation and mode-of-action studies. RT-qPCR analysis of viral genome replication in drug treated fibroblasts at 48 hpi with indicated arboviruses. Student T-test. *P >0.01, **P >0.001, ***P >0.0001. (B) (A) Dose-response curve of compounds showing antiviral activity against CHIKV. IC_50_ and R^2^ values are included for each compound. (C) Representative immunofluorescence images of indicated drug-treated and CHIKV infected human fibroblasts at 48 hpi are shown. Scale bar=25 µm. (D) Western blot analysis of innate immune pathway of cells stimulated with various STING agonists, with or without CHIKV infection at 24 hpi. Dectin-1 agonist, scleroglucan is included as an additional control.

### Comparative Transcriptomics Study on Arboviruses Treated with STING Agonist

Based on our validation data, we selected the synthetic CDN cAIMP for further mode of action studies in the context of arboviral infections using a systems-level transcriptomics approach. To uncover the dysregulated pathways in HFF-1 cells infected with these pandemic potential RNA viruses - CHIKV, WNV and ZIKV (MOI of 0.1), we performed total RNA sequencing analysis of vehicle- or cAIMP-treated virus-infected HFF-1 cells. The cells were pre-treated with the vehicle (saline), cAIMP (100 µg/ml) or scleroglucan (50 µg/ml) and, 24 hours later, viral infection was carried out. Subsequently, at 24 hpi, the cells were harvested for RNA sequencing analysis (Illumina NGS), revealing that the transcriptional responses in CHIKV-infected cells were significantly different from the host response in WNV and ZIKV (Figure 3A-B). The expression of host genes in CHIKV-infected cells were increased about 96 and 33 folds compared to WNV- and ZIKV-infected cells, respectively, reflecting differential nature of pathogenesis mechanism of these viruses and associated host responses. Because scleroglucan-treated/viral-infected cells had only a few differentially regulated genes, we mainly focused on cAIMP-treated samples for further analysis. While similar number of host genes were expressed in either vehicle- or cAIMP-treated CHIKV-infected cells, distinctive patterns of transcription was observed (Figure 3A). However, host gene expressions were increased to 32 and 6 folds in WNV- and ZIKV-infected cells treated with cAIMP compared to the vehicle-treated infected cells, respectively. This demonstrates a uniform antiviral response of cAIMP in host response to different viral infections. Considering the different levels of host transcriptional response to these vehicle- and cAIMP-treated viral infections, only 3 (*OAS2, IFI44L*, and *HELZ2*) and 74 genes, respectively, were commonly upregulated (Figure 3B). These 74 common genes could potentially be drug-specific elements contributing to the broad-spectrum antiviral process (Supplementary Table 3). Among cAIMP-treated/infected cells, the majority (88%) of the upregulated genes in ZIKV-infected cells were also upregulated in WNV-infected cells, suggesting that some specific molecular functions are commonly pursued by these two flaviviruses compared to the alphavirus CHIKV.

**Figure 3.**
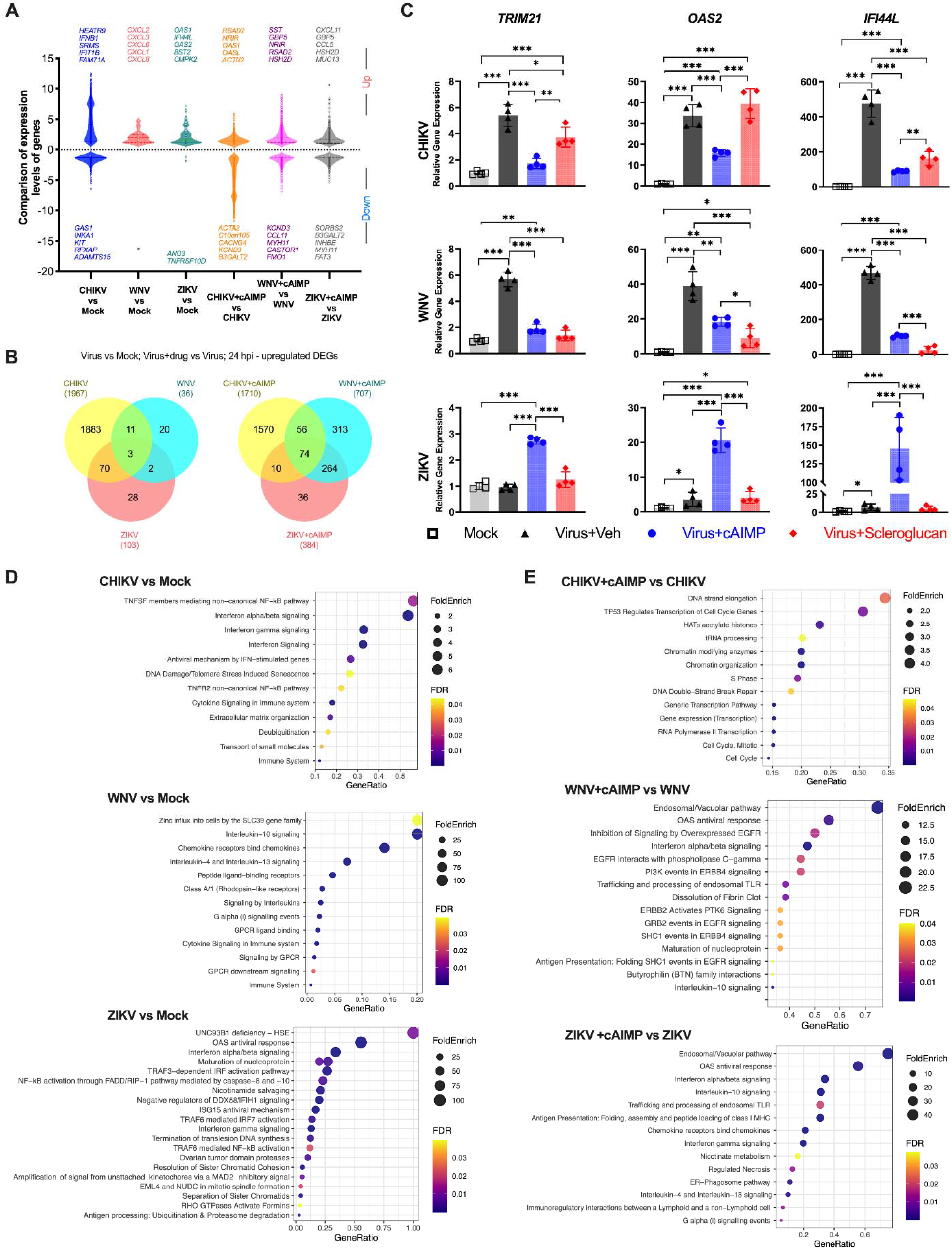
Comparative transcriptome analysis of drug treated and arboviruses infected human fibroblasts. (A) Violin plot shows comparative analysis of differentially-expressed genes in with or without cAIMP drug treated cells at 24 hpi. (B) Venn diagrams display the number of common and unique genes upregulated in virus infected cells (left) as well as in cAIMP treated virus infected cells (right) at 24 hpi (FDR < 0.01 and Log2FC>1). (C) Graphs show RT-qPCR analysis of innate immune STING pathway genes in arboviruses’ infected fibroblasts at 48 hpi with or without drug treatments of infected cells. (D-E) Dot plot analysis of overrepresented pathways in viral infected (D) and cAIMP-treated virus-infected cells (E). The size of the dot represents the fold enrichment (FoldEnrich), while its color represents the FDR (p-adjusted) value for each enriched Reactome pathway. The x-axis represents the percentage of up-regulated genes in the selected pathway that is presented in the y-axis.

We performed a sensitive RT-qPCR analysis to quantitatively assess the levels of each target gene in the STING signaling pathway following infection and treatment with drug compounds (Figure 3C; Supplementary Figure 3A). This data revealed that CHIKV and WNV activated STING pathway genes, including *cGAS, IFI16* and *TRIM21* at 48 hpi. Moreover, all three arboviruses upregulated type I interferon-stimulated genes *OAS2* and *IFI44L*. Both these STING and type 1-IFN pathway genes were transcriptionally induced in cAIMP-treated/viral-infected cells. In general, many of these innate immune pathway gene expressions were reduced by treatments with either cAIMP or scleroglucan, which could be due to inhibition of virus replication in treated cells (Figure 3C and Supplementary Figure 3A). Furthermore, we have provided the results of the expression patterns of STING pathway genes from the comparative transcriptome dataset for these arboviruses at an early timepoint of 24 hpi (Supplementary Figure 3B). Many STING pathway genes were differentially regulated in CHIKV-infected cells compared to ZIKV- or WNV-infected cells. Furthermore, in response to these virus infections, the host cells have upregulated various pathways, including IFN signaling pathways, NF-κB and immune cytokine signaling pathways, as well as virus-specific pathways, such as DNA damage/telomere stress-induced senescence (CHIKV), zinc influx into cells by the SLC39 gene family (WNV), and UNC93B1 deficiency – herpes simplex virus type 1 encephalitis (HSE) (ZIKV) (Figure 3D and Supplementary Table 2). While cAIMP-treated/virus-infected cells primarily triggered many antiviral immune responses, we observed unique upregulation of genes in the cell cycle and transcription pathways in cAIMP-treated/CHIKV-infected cells (Figure 3E and Supplementary Table 2). A complete list of pathways enriched during virus infections in the context of drug treatment is provided in the Supplementary Table 2. These results indicate that CHIKV contributes to robust transcriptional dysregulation in fibroblasts compared to WNV and ZIKV, reflecting a possible difference in the disease pathogenesis mechanisms by alphaviruses and flaviviruses despite being transmitted by the same mosquito vectors. Irrespective of the observed differences in molecular signatures, the STING agonist provided a broader protection against these arboviruses.

### Pharmacogenomics Study on STING and Dectin-1 Agonists Against CHIKV

We further performed an in-depth transcriptomics study to compare the antiviral efficacy of synthetic CDN cAIMP and scleroglucan in CHIKV-infected HFF-1 cells. To uncover the dysregulated pathways in CHIKV-infected (vehicle-treated) as well as drug-treated infected cells, we performed total RNA sequencing analysis at 24 hpi. The transcriptome data reveals that viral genome reads comprised of 83% of the total reads for vehicle-treated cells, 63% for scleroglucan-treated cells, and 0.1% for cAIMP-treated cells, in concurrence with the observed antiviral phenotype (Figure 4A). Viral count analysis showed uniform increases in expression of all viral genes (structural and non-structural) in CHIKV-infected cells. However, viral gene expression was reduced about 880 folds in cAIMP-treated infected cells compared to untreated infected cells (Figure 4B). RNA sequencing analyses indicated that the transcriptional response in CHIKV-infected cells is significantly different from the host response to the agonists, especially for cAIMP, which further explains their extreme coordinates on the principal-component analysis (PCA) plot (Figure 4C). Moreover, treatment with cAIMP in uninfected and infected cells show very similar transcriptional responses, though different from scleroglucan-treated cells (Figure 4A-E). In vehicle-treated cells, viral RNA approached 83% of total RNA reads at 24 hpi, which clusters apart from drug-treated RNA samples, reflecting an overall difference in host response (Figure 4C-E). With regard to different levels of CHIKV gene expression in treated infected cells (Figure 4B), the host transcriptional response to CHIKV infection and cAIMP-treatment CHIKV infection are differential in nature (Figure 4E) with only 39 and 230 shared down- and upregulated genes, respectively (Figure 4G), suggesting that genes involved in only certain molecular functions are commonly shared by two conditions. However, cAIMP-treated CHIKV-infected cells had 284 unique differentially expressed genes that could potentially be drug-specific elements contributing to the antiviral process. Pathway and Gene Ontology enrichment analyses of virus-infected cells treated with cAIMP showed upregulation of many cell repair mechanisms, including histone acetyltransferase (HAT) activity, DNA double-strand break repair, and metabolic pathways of amino acids and their derivatives (Supplementary Figure 4). Additionally, cAIMP-treated (uninfected cells) exhibited upregulation of multiple antiviral and metabolic pathways, including negative regulation of DDX58/IFIH1 signaling and nicotinate metabolism, which are downregulated in CHIKV-infected cells (Figure 4H and Supplementary Figure 4). Analysis of differentially regulated genes in various experimental conditions showed that HEAT Repeat Containing 9 (*HEATR9*) is highly upregulated in CHIKV-infected cells (Figure 4F) and cAIMP treatment reversed the induction of this gene. The HEATR9 gene has been shown to regulate cytokine production^32^ with mutations reported in patients with POEMS syndrome, a plasma cell dyscrasia.

**Figure 4.**
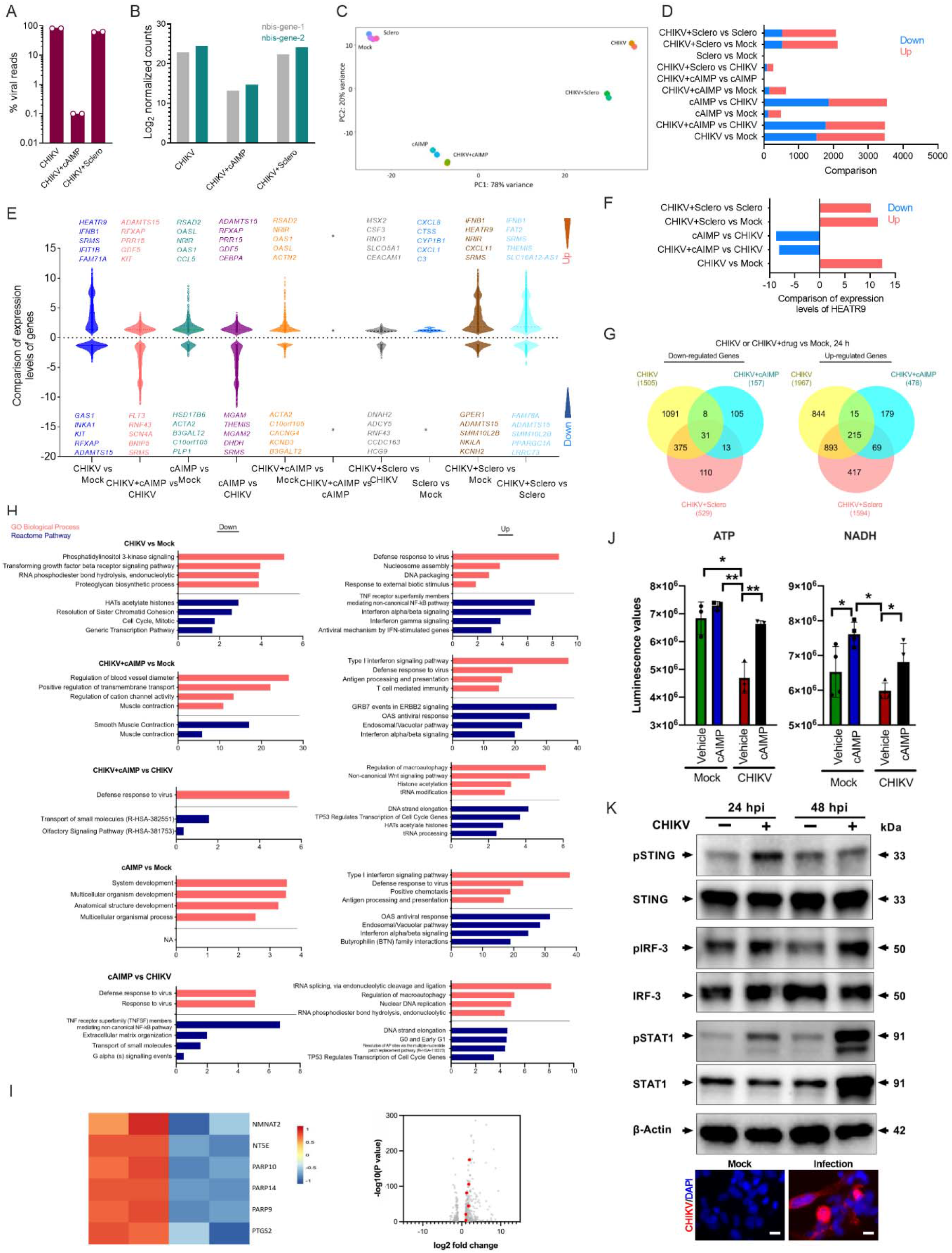
Transcriptome analysis of control and drug treated/CHIKV-infected human fibroblasts. (A) Bar graph shows proportion of total reads comprising CHIKV transcripts in indicated treatments. The proportion of virus-aligned reads over total reads is shown for each sample. Error bars represent average (± SD) from two biological replicates. (B) Normalized read counts (log2) of CHIKV RNA products, showing transcriptional enrichment of viral genes in CHIKV infected cells when compared with uninfected cells. The infected cells treated with cAIMP drug showed much less viral enrichment. (C) Principal-component analysis for the global transcriptional response to CHIKV infection and drug cAIMP or Scleroglucan treatment. (D) Identification of the number of down- and up-regulated genes in different samples. (E) Comparison of expression levels of differentially-expressed genes in different samples. The greater number of genes up-regulated in CHIKV infected cells, while greater number of genes down-regulated in cAIMP treated/CHIKV infected cells. (F) Comparison of expression levels of HEATR9 in different samples. (G) Venn diagram displays the number of common and unique genes down- and up-regulated in various conditions. CHIKV infected cells had greater numbers of down- and up-regulated genes. The CHIKV infected cells treated with cAIMP had the least number of genes down- and up-regulated. (H) In silico functional annotation of differentially expressed gene sets in different samples performed using PANTHER, and the four most overrepresented GO Biological Process (orange) and Reactome Pathway (blue) terms are shown. (I) Heatmap and Volcano plot (red; FDR < 0.01 and Log2FC>1) of upregulated genes in nicotinate metabolic pathway in cAIMP treated cells. Heatmap illustrates Z scores as expression levels of these 6 DEGs. Red and blue colors represent genes upregulated in cAIMP treated alone cells and downregulated genes in control cells, respectively. (J) Graphs show intracellular ATP and NADH metabolites of uninfected and CHIKV infected cells, with or without cAIMP treatment at 48 hpi. Student T-test. *P >0.01, **P >0.001. n=2 independent experiments. (K) Western blot analysis of STING-IRF3 innate immune pathway activation in CHIKV infected human RD muscle cell line. Immunofluorescence images show CHIKV infection in fibroblasts at 48 hpi.

In response to CHIKV infection and cell injury, the infected cells upregulated various pathways including TNF receptor superfamily-mediated NF-κB pathway, DNA-damage/telomere stress induced senescence, and immune cytokine and type I IFN signaling pathways (Figure 3D, 4H and Supplementary Table 2). However, cAIMP treatment primarily activated antiviral innate immune responses (Supplementary Figure 4). Interestingly, cAIMP treatment resulted in upregulation of nicotinate metabolic pathway (Figure 4I). Functional validation by intracellular metabolite analysis confirmed that cAIMP treatment increased nicotinamide adenine dinucleotide (NADH) level (Figure 4J). NADH can help provide energy to execute the antiviral responses. Similarity, cAIMP treatment prevented ATP depletion likely by inhibiting virus replication. We have observed phosphorylation of STING during CHIKV infection, suggesting activation of STING pathway as well as downstream IRF3-IFN-STAT1 pathway in human (RD) muscle cells (Figure 4K). Taken together, our comprehensive transcription analysis and validation revealed specific mode of mechanism of action for cAIMP treatment and pathophysiological molecular changes during CHIKV infection.

### Antiviral Compounds Prevent Viral Infection of Cardiomyocytes

Cardiovascular involvement has been shown to be a common manifestation of CHIKV infection^6^. In addition, other viral-mediated cardiovascular diseases among RNA viral infections have been reported. Human patients have been shown to develop myocarditis and pericarditis after subsequent WNV and enterovirus D68 (EV-D68) infections^33,34^. Recently, it has been shown that SARS-CoV-2 is responsible for multiple cardiovascular (CV) manifestations^35-37^ and can infect human cardiomyocytes^38,39^. Therefore, assessing a broad spectrum of antivirals in a relevant cardiomyocyte system can help determine the effectiveness of these compounds. We first evaluated the susceptibility of cardiomyocytes to viral infections using a human pluripotent stem-cell-derived cardiomyocyte (PSC-CM) system (Figure 5A). We have previously shown that SARS-CoV-2 can establish active infection in PSC-CMs^38,40^. We observed that CHIKV established productive infection in PSC-CMs, which was inhibited by scleroglucan treatment (Supplementary Figure 5A). For detailed study, we also included additional respiratory pathogens, namely respiratory syncytial virus (RSV) and EV-D68. The PSC-CMs were infected with various viruses. At 48 hpi, the cells were fixed with 4% paraformaldehyde and immunostained with antibodies targeting each virus-specific antigen (Supplementary Table 4). IHC analysis indicated that the cardiomyocytes are highly susceptible to CHIKV, WNV and EV-D68 infections, but not RSV (Figure 5A). Using this platform, we subsequently tested STING agonists cAIMP and diABZI, as well as IFN-β and remdesivir, against these arbo- and respiratory viruses (Figure 5B). The STING agonists demonstrated potent antiviral activity across all tested RNA viruses in the biologically relevant human PSC-CMs. These agonists induced phosphorylation of STING in PSC-CMs indicating the activation of STING-mediated antiviral signaling cascade (Supplementary Figure 5A). Interestingly, the STING agonists were not effective against RSV in human A549 lung epithelial cells. However, IFN-β, remdesivir, and 6-Azauridine demonstrated efficient inhibition of RSV infections (Figure 5C and Supplementary Figure 5C). Remdesivir, a well-known RNA-dependent RNA polymerase inhibitor approved for SARS-CoV-2 treatment, had no effective antiviral activity against CHIKV or WNV. Taken together, the STING agonists exhibited broad-spectrum antiviral activity against both arbo- and respiratory viruses.

**Figure 5.**
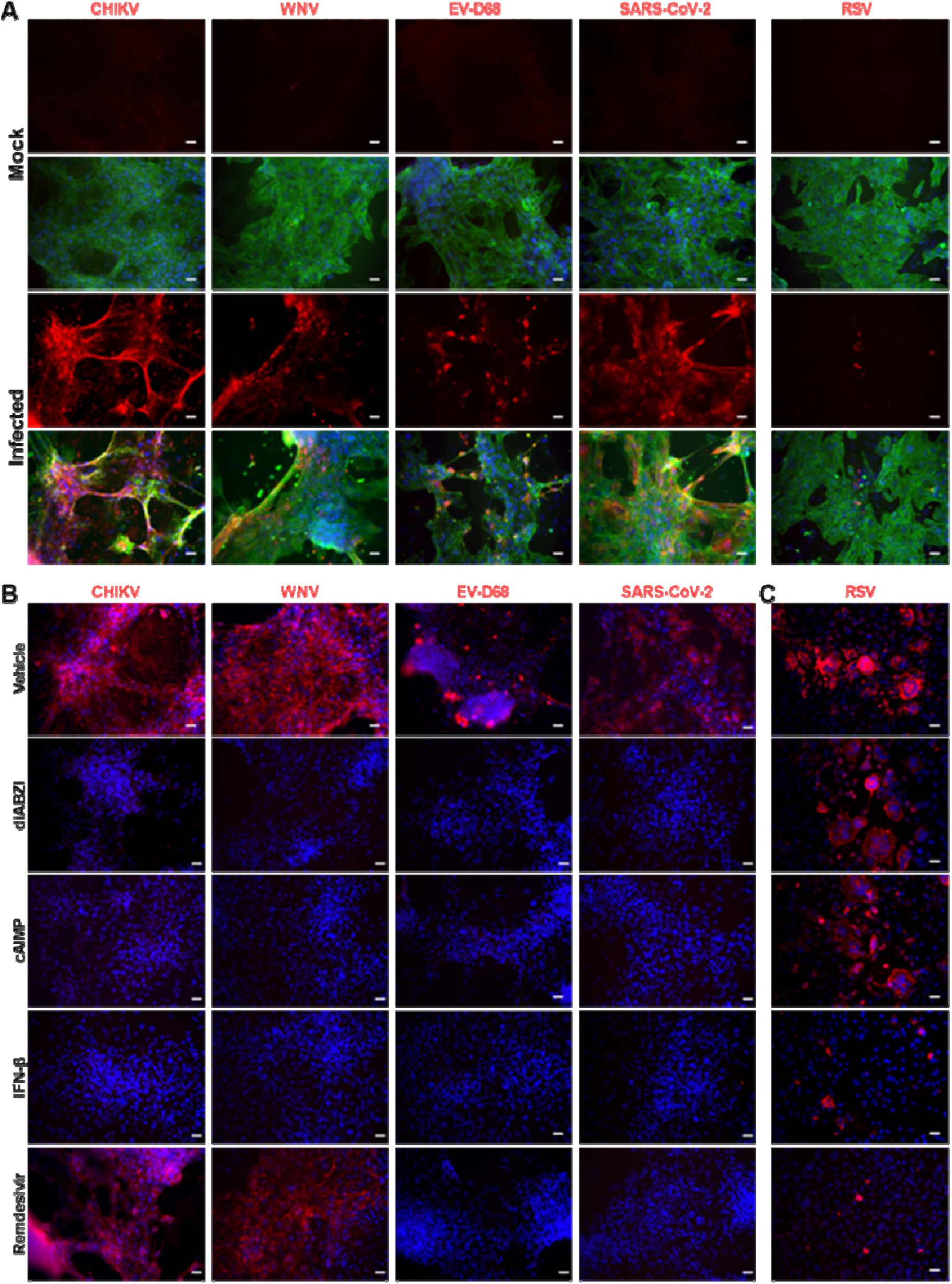
Evaluating the susceptibility of PSC-CMs to different families of RNA viruses and testing broad-spectrum antiviral therapeutic efficacy. (A) IHC images depict infected PSC-CMs with indicated RNA viruses, EV-D68 (anti-VP1 protein), SARS-CoV-2 (anti-Spike protein), and RSV strain A2001/3-12 (anti-Fusion protein), and cardiac-troponin I (green). Scale bar=25. Note: RSV is not efficient in infecting cardiomyocytes. (B) IHC images present antiviral activity of tested compounds. STING agonists and IFN-β have potent antiviral activity across all tested RNA viruses. Remdesivir is not active against CHIKV and WNV. Scale bar=25. (C) Immunofluorescent images present RSV infected AF49 lung epithelial cells at 48 hpi. Note: IFN-β and Remdesivir demonstrated efficient inhibition of RSV infection. Scale bar=25. Representative data from n=2 independent experiments are provided.

### cAIMP Provides Therapeutic Benefit in Treating CHIKV Arthritis in Mouse Model

Next, we evaluated the antiviral effect of cAIMP, a novel synthetic CDN, in a pre-clinical mouse model of CHIKV. This potent antiviral compound cAIMP, an analog of natural 3’3’-cGAMP, is derived with adenine and inosine nucleosides. The schematic of both prophylactic and therapeutic animal study designs is provided in Figure 6A. For prophylactic study, the mice (14-month-old, C57BL/6J) were systemically pretreated with a single dose of cAIMP (10 mg/kg) by intraperitoneal injection at Day -1. The control group received a saline injection as a placebo treatment. 24 hours later, a group of cAIMP or saline administered animals (n=5) were sacrificed and the left footpad tissues were harvested for gene expression analysis. The cAIMP-treated animals had significant activation of STING pathway and antiviral genes, *Trim21* and *Oas1* (Figure 6B). In parallel at 24 hours post-treatment (hpi), additional groups of mice were inoculated with CHIKV (strain 181/25) in the left rear footpad by subcutaneous injection. The animals were observed for the next 7 days for body weight changes and clinical signs of footpad swelling (Figure 6C and Supplementary Figure 6A). CHIKV infection did not have a deleterious effect on overall animal health as the infected animals maintained body weight throughout the study. We observed that a single dose of prophylactic cAIMP treatment significantly prevented CHIKV-mediated footpad swelling, whereas the thickness of viral inoculated left-rear footpad was increased in the saline-treated animals (Figure 6C). During the acute phase of infection on Day 3, mice were sacrificed and the viral titer in the inoculated left-rear footpad was measured by RT-qPCR (Figure 6C). We have observed that the cAIMP pre-treatment resulted in 2 log reduction in viral genome replication. Transcriptomics analysis of left-rear footpad at 7 dpi revealed that transcriptional differences in untreated (648 DEGs) and cAIMP-treated (1,183 DEGs) CHIKV-infected animals (Figure 6D and Supplementary Figure 6D) – a similar pattern as was observed in our analysis of human HFF-1 cells. The mouse transcriptional responses were increased to almost 2 folds in cAIMP-treated CHIKV-infected animal compared to the saline-treated CHIKV-infected animal. This indicates a robust antiviral host response induced by cAIMP in CHIKV-infected footpad. Transcriptome analysis revealed that CHIKV infection activated several biological pathways involved in interferon gamma signaling, antimicrobial proteins and platelet adhesion. Compared to the saline-treated/CHIKV-infected mice group, cAIMP-treated mice showed upregulation of myogenesis, and metabolic pathways involved in keratan sulfate, chondroitin sulfate and angiotensin conversion; meanwhile down regulation of NLRP3 inflammasome, pyroptosis and various interleukin signaling pathways was observed (Figure 6D and Supplementary Table 2). Histopathological analysis revealed that the saline group had heavy infiltration of inflammatory cells in the footpad muscle, skin, and joint tissues, resulted in myocytis, dermatitis, and arthritis at 7 dpi (Figure 6E). The synovial cavity of the affected joint had fibrinous exudate. In addition, the synovial membrane lining the joint synovial cavity was heavily infiltrated with mononuclear inflammatory cells (Figure 6E and Supplementary Figure 6B). Furthermore, IHC analysis revealed that the saline-treated/CHIKV-infected left footpad had heavy infiltration of mononuclear cells including macrophages, CD4+ helper T cells, and CD8+ cytotoxic T cells. The observed pathological changes and left-rear footpad swelling were prevented upon cAIMP administration. These gross and histopathological results were supported by significant downregulation of the expression of inflammatory genes *Ccl2, Il-6*, and *Il-10* in the left footpad of the cAIMP pre-treated group at 7 dpi (Supplementary Figure 6C). The contralateral right-rear footpad volumes were not changed respective to the inoculated left-rear footpad, suggesting the infection is predominantly localized (Supplementary Figure 6E). In a chronic long-term follow-up study up to 2 months post-infection using 14-month-old animals, cAIMP pretreatment has significantly reduced the volume of footpad swelling as well as virus replication (Supplementary Figure 6F). This long-term study showed that CHIKV can establish a chronic persistent infection in the older mice. These observations indicate that STING pathway induction exerts a strong antiviral response *in vivo*.

**Figure 6.**
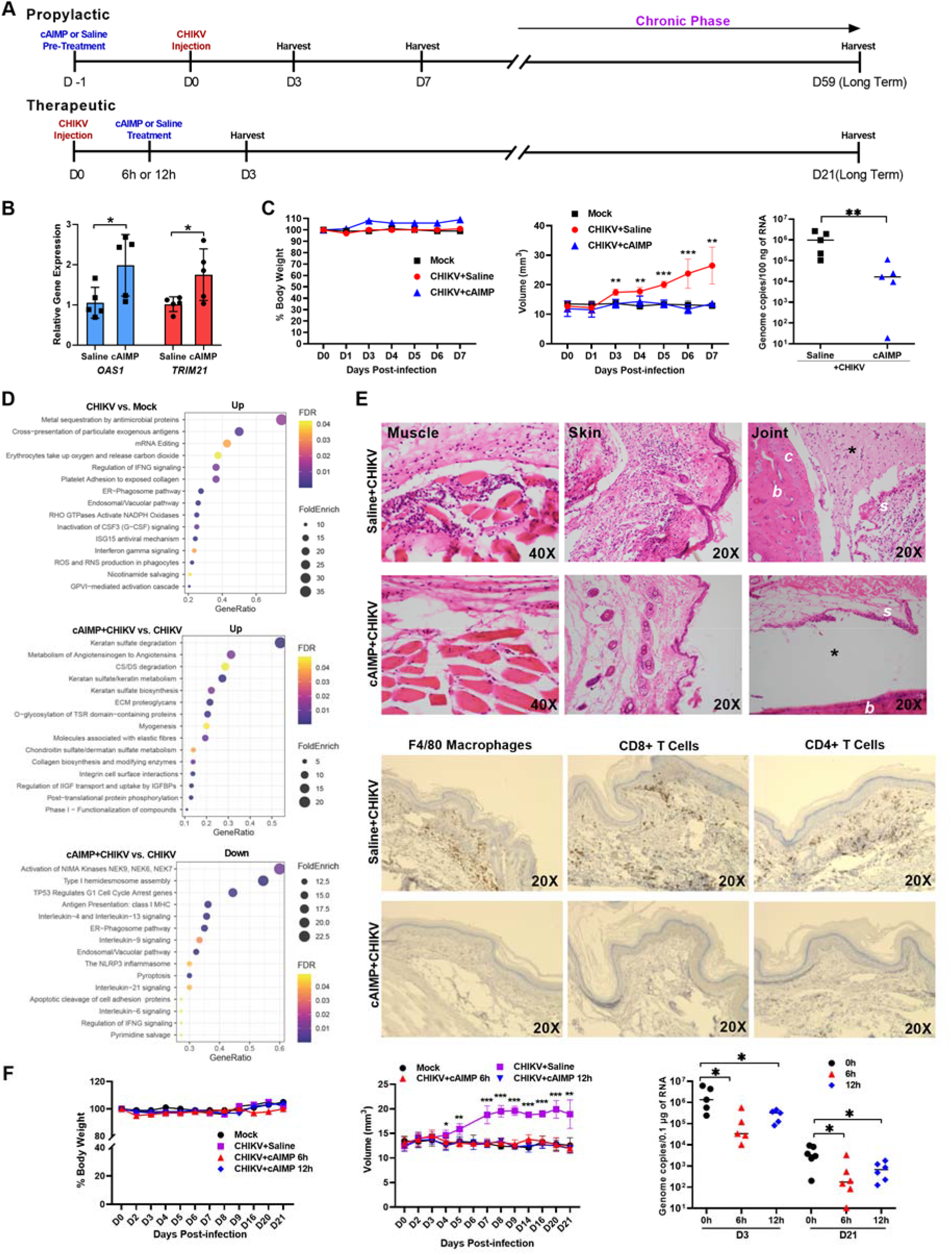
STING agonist cAIMP prophylactic and treatment measures mitigate CHIKV-mediated viral arthritis. (A) Schematic diagram of timeline highlighting time of infection, administration of the compound, and tissue harvest. (B) Systemic administration of cAIMP induces the expression of antiviral genes. Graph shows the transcriptional activation of STING/type 1 IFN genes in the left footpad at 24 h after drug treatment (n=5 mice/group). Student T-test. *P >0.01. (C) Graphs show body weight, left footpad volume and viral genome replication (left footpad at 3 dpi) of cAIMP-treated or vehicle-treated mice. cAIMP pretreatment reduces foot swelling in CHIKV (181/25) infected mice. Student T-test and a non-parametric t-test (Mann-Whitney Test) was used. **P >0.001, ***P >0.0001. n=2 independent experiments. (D) Dot plot analysis of overrepresented up or downregulated pathways in CHIKV infected, and cAIMP treated/virus infected left footpad of mice at 7 dpi are shown. (E) Histopathological analysis of CHIKV-infected left rear footpad at 7 dpi. Microscopic images of H&E staining of muscle, skin, and joint tissues are presented. Note: Heavy inflammatory cell infiltration in the saline group. Saline group synovial cavity (asterisk) in the joint is filled with fibrinous exudate. C=cartilage; B=bone; S=synovial membrane. IHC images indicate infiltrating macrophages and T cells (dark brown) in the left footpad of vehicle (saline) or cAIMP treated mice at 7 dpi. Image magnification with 20x or 40x objective lens. (F) cAIMP drug treatment at 6 or 12 hours post-CHIKV infection provides long-term therapeutic benefit by reducing footpad swelling CHIKV viral load (n=5-6 mice). No significant body weight changes observed in any of the groups. Student T-test. *P >0.01, **P >0.001, ***P >0.0001.

Subsequently, we evaluated the efficacy of cAIMP in a therapeutic capacity. The mice were infected with CHIKV in the left-rear footpad and treated with a single dose of cAIMP (10 mg/kg) either at 6 hpi or 12 hpi via systemic intraperitoneal route (Figure 6F). The animals were observed up to 21 dpi for clinical signs of footpad swelling and body weight change. Similar to the prophylactic countermeasure, we noticed that the cAIMP treatment has protected the animals from significant footpad swelling by reducing viral replication (Figure 6F), reducing inflammatory gene expression and inflammatory cellular infiltration (Supplementary Figure 7). Taken together, the STING agonist cAIMP has demonstrated potent prophylactic and therapeutic efficacy against CHIKV arthritis.

Based on our *in vitro* and *in vivo* results, we have provided a schematic illustration of our hypothetical model of the tested innate immune agonists displaying broad-spectrum antiviral activities (Supplementary Figure 8). The availability of these PRR agonists for antiviral therapy could prove critical for future use against these pandemic potential viruses.

## DISCUSSION

The objective of this study was to identify broadly acting antivirals against pandemic potential arthropod-borne viruses. Importantly, we observed that agonists of the cGAS-STING cytosolic DNA sensing pathway exhibited potent antiviral activity against members of multiple sense strand RNA viral families. STING is a core component of the cytosolic DNA sensing pathway. However, the mechanism of STING pathway-mediated antiviral response against RNA viruses is not clear. STING can be involved in sensing RNA viruses through cross-talk with the RNA-sensing RIG-I-MAVS pathway and mitochondria DNA leakage in infected cells^41-47^. In *Drosophila melanogaster*, cGAS-like receptors have been described for sensing RNA sequences and activating STING pathway^48^. There may have been cGAS-like receptors in mammalian cells that can recognize viral RNA. Deficiency of cGAS and STING in the embryonic fibroblasts of golden ticket mice (I199N missense mutation in *Sting* resulting in lack of IFN-β production following STING agonist stimulation) saw significantly exacerbated infection by CHIKV, suggesting that the cGAS-STING pathway may play a role in attenuation of Chikungunya disease. This finding supports previous work demonstrating its role in the restriction of alphavirus replication^49-52^. The CHIKV nsP1 protein, a non-structural protein of CHIKV which anchors to cell membranes, has also been found to interact directly with STING. This interaction elucidates an important immune evasion role of nsP1 in disrupting STING dimerization and resultant dampening of STING-mediated immune response to CHIKV infection^49,53-55^. The CHIKV capsid protein was found to induce autophagy-dependent degradation of cGAS, leading to downregulation of IFN-β transcription^56^. Moreover, a previous study has shown that Sindbis virus nsP4 is degraded by the proteosome^57^. In this context, observed interactions between CHIKV nsP4 and cGAS^49^ may lead to partial cGAS degradation via the proteosome. However, there is no direct evidence to support this. Beyond demonstrating the immune-dampening potential of CHIKV, this observation reveals a direct evolutionary mechanism within the RNA virus that targets the cGAS-STING pathway, thus emphasizing the need to better understand the role of cGAS-STING in the modulation of CHIKV-induced arthritis and myositis. We have observed that stimulation of this pathway with both natural and synthetic STING-activating compounds overcomes CHIKV-mediated immune evasive mechanisms. Moreover, STING agonists have been shown to exert antiviral activity against SARS-CoV-2^58^. In addition, evidence suggests that STING-activating compounds can be useful in mediating a robust immune response as adjuvants in vaccines against viral infections (HIV, influenza, and coronaviruses) and can be used as initiators of anticancer immunostimulatory effects^59,60^. This unique avenue for constructing effective vaccines is possible due to stimulation of the cGAS-STING pathway and activation thereafter of IRF3, NF-κb, type I IFNs, and other pro-inflammatory cytokines that help modulate antigen presentation and immune responses^59,61-64^.

In comparing the genome replication and transcriptome of CHIKV with other viruses, we observed a higher number of gene dysregulation and a higher viral load. This finding suggests that CHIKV displays faster replication kinetics in our *in vitro* models than WNV or ZIKV. Even in cases of co-infection with other viruses, such as DENV and ZIKV, CHIKV replication has been found to either not affected^65^ or results in decreased replication^66^ of the co-infecting virus. Our finding has been substantiated in other studies^65,67,68^, suggesting a ready path to widespread infection and greater pandemic potential for CHIKV.

Interestingly, we observed that the Dectin-1 ligand scleroglucan exhibited antiviral activity by preventing infection of arboviruses in fibroblasts, as well as moderately reducing CHIKV replication in PSC-CMs. The mode of action of this activity is independent of the STING-IRF3-STAT1 type I IFN pathway. Transcriptomics analysis confirmed that scleroglucan can only stimulate the inflammatory pathway. Compared to the STING agonist, cAIMP activated the type I IFN signaling pathway and defense response against the virus. Additionally, gene knockdown experiments can provide valuable insight into the key genes involved in anti-viral response triggered by these compounds against the tested RNA viruses. We have noticed that the compound Pam3CSK4, a synthetic triacylated lipopeptide^55^ ligand of TLR2/TLR1, had a dose-dependent enhancement of cell death in CHIKV-infected cells, although the compound alone has no toxicity. This observation indicates that, in this cell type, CHIKV infection can be aggravated in the presence of bacterial cell wall components. Thus, it is possible that comorbidities, particularly those attributed to bacterial infections, can stimulate a signaling crosstalk that might trigger apoptosis and/or cell injury.

Moreover, we have provided evidence that CHIKV, WNV, and EV-D68 pathogens can directly infect human heart cells. This reveals an important discovery of a biologically relevant platform to understand the pathogenic mechanism of cardiovascular complications caused by these viruses and to evaluate additional antiviral agents. The tested negative-strand RNA virus RSV has not shown tropism for cardiomyocytes and was not inhibited by STING agonists in transformed lung cells. Despite RSV having some association with sinoatrial blocks and transient rhythm alterations^69^, there was no direct infection of PSC-CMs. Thus, it is possible that the cardiovascular effect could be a direct outcome of RSV pulmonary infection. Additional studies are required to delineate these heart tropisms and STING pathway interactions with RSV.

In this study, we have only verified a single-dose regimen for the *in vivo* efficacy against CHIKV in a prophylactic and therapeutic setting. Further studies are required to evaluate a multiple-dose regiment of the STING agonists and additional compounds across multiple RNA viruses in animal models to optimize treatment conditions. The drug formulations can be developed for oral, inhalational, and topical modes of treatment. We have demonstrated in a CHIKV *in vivo* model that there is a long-term viral persistence for months. Therefore, further studies are required to evaluate a multimodal treatment of using STING agonists during chronic persistent arthritic phase to reduce viral induced inflammation and eliminate persistent infection. The direct-acting antiviral (DAA) remdesivir exhibited selectivity against the tested viruses. It has been demonstrated that remdesivir is readily incorporated by SARS-CoV-2 RNA-dependent RNA polymerase (RdRp) into the newly synthesized RNA strand, resulting in RdRp stalling^70^. This mechanism is more likely to be conserved in EV-D68 and RSV, whereas remdesivir was not effective against CHIKV and WNV, possibly due to evolutionary divergence of mosquito-borne arboviral RdRp structure. Thus, targeting common host factors can provide broader protection. The compounds identified in this study can be further developed and made readily available in the event of future respiratory and arboviral disease outbreaks.

## Acknowledgments

We are grateful to Barbara Dillon, UCLA High Containment Program Director for BSL3 work. We thank Yijie Wang from the UCLA Cardiomyocyte Core for providing hPSC-CM. This study is partly supported by National Institute of Health awards 1R01EY032149-01, 5R01AI163216-02 and 1R01DK132735-01 to VA. AR is supported by the Tata Institute for Genetics and Society. The viruses used in this study were obtained through BEI Resources, NIAID, NIH.

## Author contributions

Garcia Jr. G: Conception and design, Collection and/or assembly of data. Data analysis and interpretation, and Manuscript writing.

Ignatius Irudayam J., Jeyachandran AV., Dubey S., Chang C., Cario SC., Price N., Arumugam S., Shah A., Marquez AL., Fanaei A.: Conducted experiments, Data analysis and interpretation. Chakravarty N., Sinha S., French SW., Joshi S., Parcells M: Experimental design, Data analysis, Data interpretation and Manuscript writing.

Ramaiah A.: Conception and design, Bioinformatics data analysis and interpretation, Manuscript writing.

Arumugaswami V: Conception and design, Data analysis and interpretation, Manuscript writing and Final approval of manuscript.

## Data and materials availability

All mentioned and relevant data regarding this study is available from the above listed authors. In addition, supplementary information is available for this paper. Correspondence and requests for data and materials should be addressed to lead contact, Vaithilingaraja Arumugaswami.

## METHODS AND MATERIALS

### Ethics Statement

This study was performed in strict accordance with the recommendations of UCLA. All WNV, CHIKV and SARS-CoV-2 live virus experiments were performed at the UCLA BSL3 High containment facility.

### Cells

HFF-1 (SCRC-1041) and Vero E6 [VERO C1008 (CRL-1586)] cells were obtained from ATCC. HFF-1 and Vero E6 cells were cultured in Dulbecco’s Modified Eagle’s Medium (DMEM) (Gibco) and Eagle’s Minimum Essential Medium (EMEM) (Corning), respectively. DMEM contained 15% fetal bovine serum (FBS) and penicillin (100 units/ml), whereas EMEM growth media contained 15% FBS and penicillin (100 units/ml). Adenocarcinomic human alveolar basal epithelial A549 (CCL-185) cells from ATCC were cultured in the F12K (ATCC) media with the presence of 10% FBS and penicillin (100 units/ml). Cells were incubated at 37°C with 5% CO_2_. Human pluripotent stem cell-derived cardiomyocytes (hPSC-CM) were provided by UCLA Cardiomyocyte Core and were derived as described below. The PSC-CMs were differentiated from hESC line H9 using a previously described method^71^. The hPSCs were maintained in mTeSR1 (STEMCELL Technology) and RPMI1640 [supplemented with B27 minus insulin (Invitrogen)] was used as differentiation medium. From Days 0-1, 6 μM CHIR99021 was added into differentiation medium. On Days 3-5, 5 μM IWR1 (Sigma-Aldrich) was added to the differentiation medium. Thereafter, on Day 7, RPMI 1640 plus B27 maintenance medium was added. Finally, on Days 10-11, RPMI 1640 without D-glucose and supplemented with B27 was transiently used for metabolic purification of CMs ^38^.

### Drug library and compounds

The compounds tested were obtained from InvivoGen, Millipore Sigma and Selleckchem (Supplementary Table 4). A selected library of PAMP molecules was procured from InvivoGen since this library contains inhibitors for many key PRRs. All compounds were provided lyophilized and reconstituted in Nuclease-Free water (Invitrogen) or DMSO at the recommended solubility by manufacturer. Lyophilized and reconstituted compounds were aliquoted and stored at -20°C. Repeated freeze-thaw circles were avoided whenever possible.

### Viruses

CHIKV, WNV, ZIKV, EV-D68, RSV, SARS-Related Coronavirus 2 (SARS-CoV-2, Isolate USA-WA1/2020), were obtained from BEI Resources of National Institute of Allergy and Infectious Diseases (NIAID) or ATCC (Supplementary Table 4). CHIKV, WNV, and SARS-CoV-2 were passaged once in Vero E6 cells and sequence verified viral stocks were aliquoted and stored at -80°C. Virus titer was measured in Vero E6 cells by established TCID50 assay.

### Viral Infection for drug testing

HFF-1 cells were seeded at 1×10^4^ cells per well in 0.2 ml volumes using a 96-well plate. Cells were treated with indicated compounds. 24 hours later, viral inoculum of CHIKV (MOI of 0.1; 100 µl/well), WNV (MOI of 1; 100 µl/well), or ZIKV (MOI of 0.1; 100 µl/well), was added onto HFF-1 cells using serum-free base media. The hPSC-CMs were plated at 1 × 10^5^ cells per well in a 48-well plate. For hPSC-CMs, in addition to the previously-mentioned viruses, 100 µl of prepared inoculum of EV-D68 (MOI of 0.1; 100 µl/well), RSV (MOI of 0.1; 100 µl/well), or SARS-CoV-2 (MOI of 0.01; 100 µl/well) was added onto cells after removing the conditioned media from each well. After 1 hour incubation at 37ºC with 5% CO_2_, inoculum was replaced with RPMI 1640 + B27 supplement with insulin. Cells were then fixed at selected timepoints with 4% PFA, collected by 1xRIPA for protein analysis, and/or supernatant collected for viral titer. Viral infection was examined by immunostaining or Western blot analysis using antigen-specific antibodies (Supplementary Table 4).

### Viral Titer by TCID50 (Median Tissue Culture Infectious Dose) assay

The method used to measure viral production by infected cells was accomplished by quantifying TCID50 as previously described^72^. Briefly, Vero E6 cells (density of 5 ×10^3^cells/well) were plated in 96-well plates. The next day, culture media samples collected from cardiomyocytes at various timepoints were subjected to 10-fold serial dilutions (10^1^ to 10^8^) and added onto Vero E6 cells. The cells were incubated at 37ºC with 5% CO_2_. Then 3 to 4 days after, each inoculated well was carefully evaluated for presence or absence of viral CPE. Thereafter, the percent infected dilutions immediately above and immediately below 50% were determined. TCID50 was calculated based on the method of Reed and Muench.

### Cell Viability and ATP Assay

We performed Cell-Titer Glo Luminescent Assay (Promega) as indicated by manufacturer for assessing viability and intracellular ATP level. HFF-1 cells were seeded on 96-well plates. After 48 hours of drug treatment, a working reagent (100 µl) was added to the cells and incubated for 30 minutes at room temperature. Thereafter, 100 µl of the cell-reagent reaction was transferred to a 96-well white bottom plate. The luminescence of each condition was measured in triplicate values and recorded. Percent viability for each compound was calculated based on vehicle (water or DMSO) treated cells.

### NAD/NADH-Glo Assay

We performed NAD/NADH Glo Luminescent Assay (Promega) as indicated by manufacturer for measuring intracellular NADH level. HFF-1 cells were seeded on 96-well plates and subjected to drug treatment and CHIKV infection. After 48 hpi, a working reagent (50 µl) was added to the cells (50 µl media volume) and incubated for 45 minutes at room temperature. The luminescence signal of 100 µl of the cell-reagent reaction was quantified in triplicate values and analyzed.

### Histopathology

Histopathological services were provided by UCLA Translational Pathology Core Lab. Mice footpad samples were processed, decalcified and sectioned for H&E staining, and subsequent image analysis. Immunohistochemistry stainings were also performed on these footpad tissues: Paraffin-embedded sections were cut at 4-μm thickness and paraffin was removed with xylene and the sections were rehydrated through graded ethanol. Endogenous peroxidase activity was blocked with 3% hydrogen peroxide in methanol for 10 minutes. Heat-induced antigen retrieval was carried out for all sections in AR9 buffer (AR9001KT Akoya) using a Biocare decloaker at 95°C for 25 minutes. The slides were then stained with primary antibodies targeting mouse F4/80, CD4 and CD8 antigens at 4ºC overnight; the signal was detected using Bond Polymer Refine Detection Kit (Leica Microsystems, catalogue #DS9800) with a diaminobenzidine reaction to detect antibody labeling and hematoxylin counterstaining.

### Immunohistochemistry

Cells were fixed with methanol (incubated in -20°C freezer until washed with PBS) or 4% paraformaldehyde for 30-60 minutes. Cells were washed 3 times with 1x PBS and permeabilized using blocking buffer (0.3% Triton X-100, 2% BSA, 5% Goat Serum, 5% Donkey Serum in 1 X PBS) for 1 hour at room temperature. For immunostaining, cells were incubated overnight at 4ºC with each primary antibody. The cells were then washed with 1X PBS three times and incubated with respective secondary antibody for 1 hour at room temperature. Nuclei were stained with DAPI (4’,6-Diamidino-2-Phenylindole, Dihydrochloride) (Life Technologies) at a dilution of 1:5000 in 1X PBS. Image acquisition was done using Leica DM IRB fluorescent microscopes.

### Image Analysis/Quantification

Microscope images were obtained using the Leica DM IL LED Fluo and Leica LAS X Software Program. Then, 4-8 images per well were taken and quantified for each condition and timepoint using Image J’s plugin Multipoint and Cell Counter feature to count the positively-stained cells by a double blinded approach.

### Viral infection and RNA sample preparation for RNA sequencing analysis

To determine cellular response to drug treatment and CHIKV infection, the HFF-1 cells in 12-well plate format were subjected to either vehicle or drug (cAIMP or Scleroglucan at 100 µg/ml) pretreatment. The next day, cells were inoculated with individual CHIKV, WNV or ZIKV (MOI of 0.1) and after 1 hour of incubation for viral adsorption, the inoculum were replaced with complete DMEM media. We included mock-infected but vehicle-treated cells as negative control. We used quadruplicate wells for each condition. 24hpi, the cells were lysed with 1 ml of Trizol and proceeded with total RNA isolation as described below. The bulk RNA was extracted using RNeasy Mini Kit (Qiagen), as per the manufacturer’s instructions. RNA was quantified using a NanoDrop 1,000 Spectrophotometer (Thermo Fisher Scientific). Duplicate RNA samples per treatment condition were submitted to the UCLA Technology Center for Genomics & Bioinformatics (TCGB) for RNA sequencing analysis.

### RNA sequencing data analysis

Libraries for RNA-Seq that were prepared with KAPA Stranded mRNA-Seq Kit. The workflow consists of mRNA enrichment and fragmentation, first strand cDNA synthesis using random priming followed by second strand synthesis converting cDNA:RNA hybrid to double-stranded cDNA (dscDNA), and incorporates dUTP into the second cDNA strand. cDNA generation is followed by end repair to generate blunt ends, A-tailing, adaptor ligation and PCR amplification. Different adaptors were used for multiplexing samples in one lane. Sequencing was performed on Illumina NovaSeq 6000 for PE 2×50 run. Data quality check was done on Illumina SAV. Demultiplexing was performed with Illumina Bcl2fastq v2.19.1.403 software. Partek Flow ^73^ as used for all data analysis. Illumina reads from all HFF-1 samples were mapped to human (GRCh38) reference genome using STAR 2.7.9a^74^ and subsequently the read counts per gene were quantified. For CHIKV data, the reads were mapped to combined human (GRCh38) and Chikungunya virus (NC_004162.2) reference genome to additionally quantify the read counts per CHIKV genes. The differential gene expression analysis was performed using DESeq2 v1.28.1 in R v4.0.3^75^ Median of ratios method was used to normalize expression counts for each gene in all samples studied. Each gene in the samples was fitted into a negative binomial generalized linear model. Genes that expressed differentially were considered only if they were supported by a false discovery rate (FDR) *p* < 0.01 and Log2 Fold Change (FC) more than 1 and -1 for up- and down-regulated genes, respectively. Unsupervised principal component analysis (PCA) was performed using DESeq2 in R v4.1.1. The gene ontology (GO) enrichment overrepresentation test was performed in PANTHER v16.0^76^ using PANTHER GO-SLIM Biological Process annotation data set^77^. Reactome pathway analysis was also performed for DEGs using human all genes as reference data set in the Reactome v65^78^ implemented in PANTHER. GO and Reactome pathway were only considered if they were supported by FDR P < 0.05. The ggplot2 v3.3.5 in R and Prism GraphPad v8.4.3 were used to generate figures. The heatmaps were generated using pheatmap v1.0.12 in R. RNA-seq data were deposited to the NCBI GEO under the accession number GSE197744.

### Western Blot analysis

For protein analysis, cells were lysed in 50 mM Tris pH 7.4, 1% NP-40, 0.25% sodium deoxycholate, 1 mM EDTA, 150 mM NaCl, 1 mM Na3VO4, 20 Mm or NaF, 1mM PMSF, 2 mg ml^-1^ aprotinin, 2 mg ml^-1^ leupeptin and 0.7 mg ml^-1^ pepstatin or Laemmli Sample Buffer (Bio Rad, Hercules, CA). Cell lysates were resolved by SDS-PAGE using 10% gradient gels (Bio-Rad) and transferred to a 0.2 µm PVDF membrane (Bio-Rad). After the transfer, the membranes were blocked (5% skim milk and 0.1% Tween-20) in 1x TBST (0.1% Tween-20) at room temperature (RT) for 1 hour. The membranes were then incubated with respective monoclonal antibodies overnight at 4ºC and detected by SuperSignal West Femto Maximum Sensitivity Substrate (Thermo Scientific). Membranes were visualized with Bio-Rad ChemiDoc MP Imaging System.

### *In vivo* mice experiment for CHIKV infection

C57BL/6J mice were used for infection study. Mice were housed at UCLA Vivarium. 14-month-old mixed sex mice (n=5-6) were prophylactically (one day before CHIKV infection) or therapeutically (6h or 12h after CHIKV infection) treated with cAIMP (10 mg/kg) by intraperitoneal injection. The control group (n=5) received only saline injection. The mice were inoculated with CHIKV (strain 181/25;1 ×10^5^ pfu per mouse in a 20 μl volume) in the left rear footpad by subcutaneous injection. The animals were observed for the next 7-59 days for clinical signs of footpad swelling. The footpad volume was measured using caliber and mice were humanely euthanized for tissue collection for histopathological analysis. Illumina reads from all animal rear left footpad RNA samples (7 dpi) were mapped to mouse (mm39) reference genome using STAR 2.7.9a^74^ and subsequently the read counts per gene were quantified. The differential gene expression analysis was performed using DESeq2 v1.28.1 in R v4.0.3^75^. Genes that expressed differentially were considered only if they were supported by a false discovery rate FDR < 0.05. Reactome pathway analysis was performed for DEGs using *Mus musculus* all genes as reference data set in the Reactome v65^78^ implemented in PANTHER. The overrepresented pathways in up or downregulated gene sets were only considered if they were supported by FDR < 0.05.

### Statistics and Data analysis

Using GraphPad Prism, version 8.1.2, IC50 values were obtained by fitting a sigmoidal curve onto the data of an eight-point dose response curve experiment. In addition, data was analyzed for statistical significance using unpaired student’s *t*-test to compare two groups (uninfected vs. infected) or a non-parametric t-test (Mann-Whitney Test) with GraphPad Prism software, also version 8.1.2 (GraphPad Software, US). All statistical testing was performed at the two-sided alpha level of 0.05.

