## Supplemental File for "Broad-spectrum antiviral inhibitors targeting pandemic potential RNA viruses"

#### SUPPLEMENTARY TABLES

**Supplementary Table 1.** List of compounds used in the primary screen and their PAMP target, cytotoxicity (percent cell viability), and respective antiviral activity at various concentrations. (Attached Separately).

**Supplementary Table 2.** Human and animal data - In silico functional annotation of differentially expressed gene sets in different conditions performed using PANTHER, and the list of overrepresented up and downregulated Reactome Pathways are shown. (Attached Separately).

**Supplementary Table 3.** List of 74 commonly upregulated genes in cAIMP treated virus infected cells potentially working as drug specific elements.

**Supplementary Table 4.** Reagents or resources used in this study. Including primer sets and list of compounds used in the secondary screen and their molecular formula, CAS number, and structure. (Attached Separately).

#### SUPPLEMENTARY FIGURE LEGENDS

**Supplementary Figure 1. Analysis of drug cytotoxicity and NF-κB phosphorylation status.** (A) Graphs show percent cell viability of selected compounds from primary screen at 72 hours post-treatment in human fibroblasts. All compounds are non-toxic at tested doses. Horizontal dotted line indicates 80% cell viability. (B) Representative bright-field images of selected compounds at 48 hours post-infection. Note: Vehicle-treated and CHIKV-infected wells had cytopathic effect (CPE). At effective concentration, the compound-treated wells did not show observable CPE. (C) Immunofluorescence images show NF-κB p65 form in fibroblast after 2-hour stimulation with indicated compounds. Scale bar=10 μm.

**Supplementary Figure 2. Drug efficacy and toxicity studies.** (A) Graphs show virus titer of vehicle or various drug treated groups at 48 hpi. Student T-test. ***P >0.0001. (B) Graph shows percent cell viability of Pam3CSK4 (TLR2/1 agonist) treated human fibroblast (72 hours post-treatment) without infection. (C) CHIKV-mediated cell death was aggravated by Pam3CSK4 treatment at a dose-dependent manner. (D) IHC images show antiviral efficacy of STING agonists against CHIKV vaccine strain (181/25; MOI 0.1) in HFF-1 cells at 48 hpi. (E) IHC images of CHIKV infected HFF-1 cells treated with cAIMP or diABZI at 2 hours post infection. An MOI of 0.1 was used and cells were fixed at 48 hpi for immunostaining analysis.

**Supplementary Figure 3.** (A) Graphs show RT-qPCR analysis of STING pathway genes in arboviruses infected fibroblasts at 48 hpi with or without drug treatments. Student T-test. *P >0.01, **P >0.001, ***P >0.0001. (B) Transcriptome analysis shows STING pathway genes that were differentially expressed in virus infected, and cAIMP treated/virus infected cells at 24 hpi. The Log2FC value of these genes (FDR <0.05) is displayed. Note: ZIKV infection alone had upregulated only two STING pathway genes. No STING pathway gene is differentially expressed for WNV infection at 24 hpi.

**Supplementary Figure 4. Pathway heatmaps and dot plot analysis.** (A-E) Heatmaps illustrate expression levels of the genes involved in indicated pathways at 24 hpi in different conditions. Red and blue colors represent upregulated and downregulated genes, respectively. Volcano plot for type 1 IFN genes upregulated in CHIKV infected cells are shown. (F) Dot plot analysis of overrepresented upregulated pathways in cAIMP alone treated cells at 24 hpi are shown.

**Supplementary Figure 5. CHIKV infection of human PSC-CMs as well as respiratory syncytial virus infection of lung cells and antiviral testing..** (A) Graph shows CHIKV infectious viral titer (TCID50) of infected PSC-CMs with or without scleroglucan treatment. Unpaired two-tailed T-test with a p value of <0.05. (B) Western blot analysis of PSC-cardiomyocytes treated with STING agonists at 3 hour timepoint. STING phosphorylation is specific to STING agonist treated cells. (C) Immunofluorescent images present RSV (strain A2000/3-4) infected and/or treated A549 lung epithelial cells. Note: 6-Azauridine and remdesivir demonstrated efficient inhibition of RSV infection and decrease of apoptosis activity (green) compared to vehicle-treated. Scale bar=25.

**Supplementary Figure 6. Short term and long term preclinical *in vivo* drug evaluation studies.** (A) Image of hind limb footpads of CHIKV (181/25)-infected mouse at 7 dpi. IL=Inoculated leg; CL=contralateral leg. Note: IL footpad is swollen compared to CL footpad. (B) Histopathological analysis of CHIKV-induced viral arthritis. Note: The synovial membrane projection (S) has heavy inflammatory cell infiltration. Similarly, subcutaneous tissue, including blood vessel (V) has inflammatory changes. C=cartilage; B=bone. (C) RT-qPCR analysis show cAIMP pretreatment result in significant downregulation of inflammatory and immune genes in the left footpad of CHIKV (181/25) infected mice at 7 dpi. Student T-test. **P >0.001. (D) Dot plot analysis of up- or down-regulated pathways in cAIMP treated/virus infected left footpad compared to that of uninfected mock at 7 dpi are presented. (E) Graphs show volume and viral load in right footpad at indicated timepoints. (F) Long term safety and efficacy study. cAIMP drug pretreatment provides long-term therapeutic benefit by reducing footpad swelling CHIKV and viral load (59 dpi) (n=6 mice/group). Student T-test. *P >0.01, **P >0.001, ***P >0.0001.

**Supplementary Figure 7.** Therapeutic evaluation of cAIMP drug in a chronic CHIKV arthritis model. (A) RT-qPCR analysis of immune and inflammatory gene expressions in the left footpad of cAIMP treated animals at 29 dpi. (B) Histopathological analysis of CHIKV-infected left rear footpad at 29 dpi. Microscopic images of H&E staining show heavy inflammatory cell infiltration in the saline group. (C) IHC images indicate infiltrating macrophages and T cells (dark brown) in the left footpad of vehicle (saline) or cAIMP treated mice at 29 dpi. Image magnification with 20x objective lens.

**Supplementary Figure 8.** Schematic illustration of hypothetical model. (C). Schematic diagram of our hypothetical model integrating innate immune agonists displaying broad-spectrum antiviral activities, which target various pathogen recognition receptors (TLRs, STING, NOD, Dectin and cytosolic DNA or RNA sensors). c-GAS, cyclic GMP-AMP synthase; IKKe, inhibitor of nuclear factor kappa B kinase subunit epsilon; MAVS, mitochondrial antiviral-signaling protein; RIG-I, retinoic acid inducible gene I protein; STING, stimulator of interferon response cGAMP interactor.

**SUPPLEMENTAL FIGURES**

**Supplementary Figure 1**

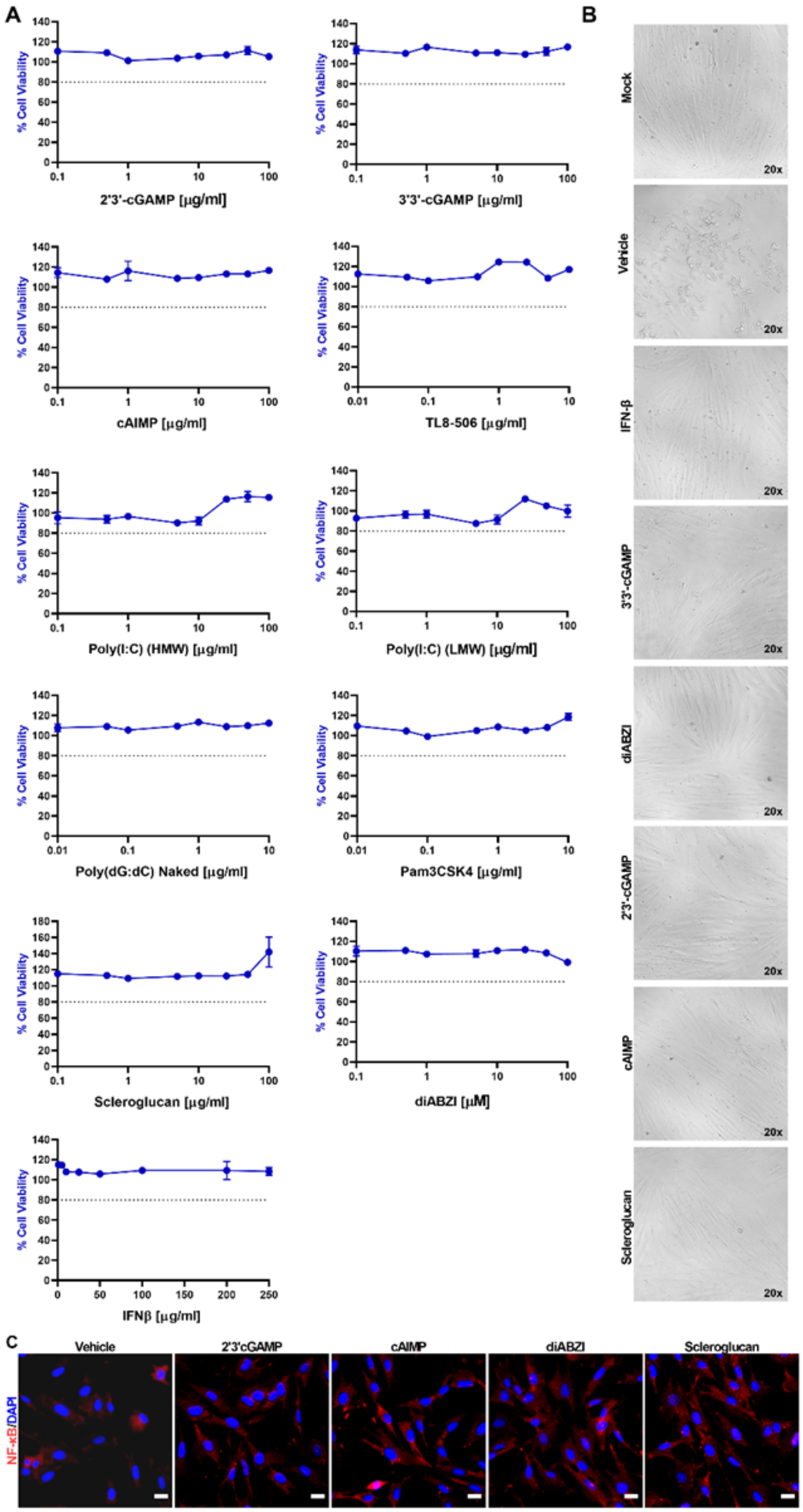

#### Supplementary Figure 2

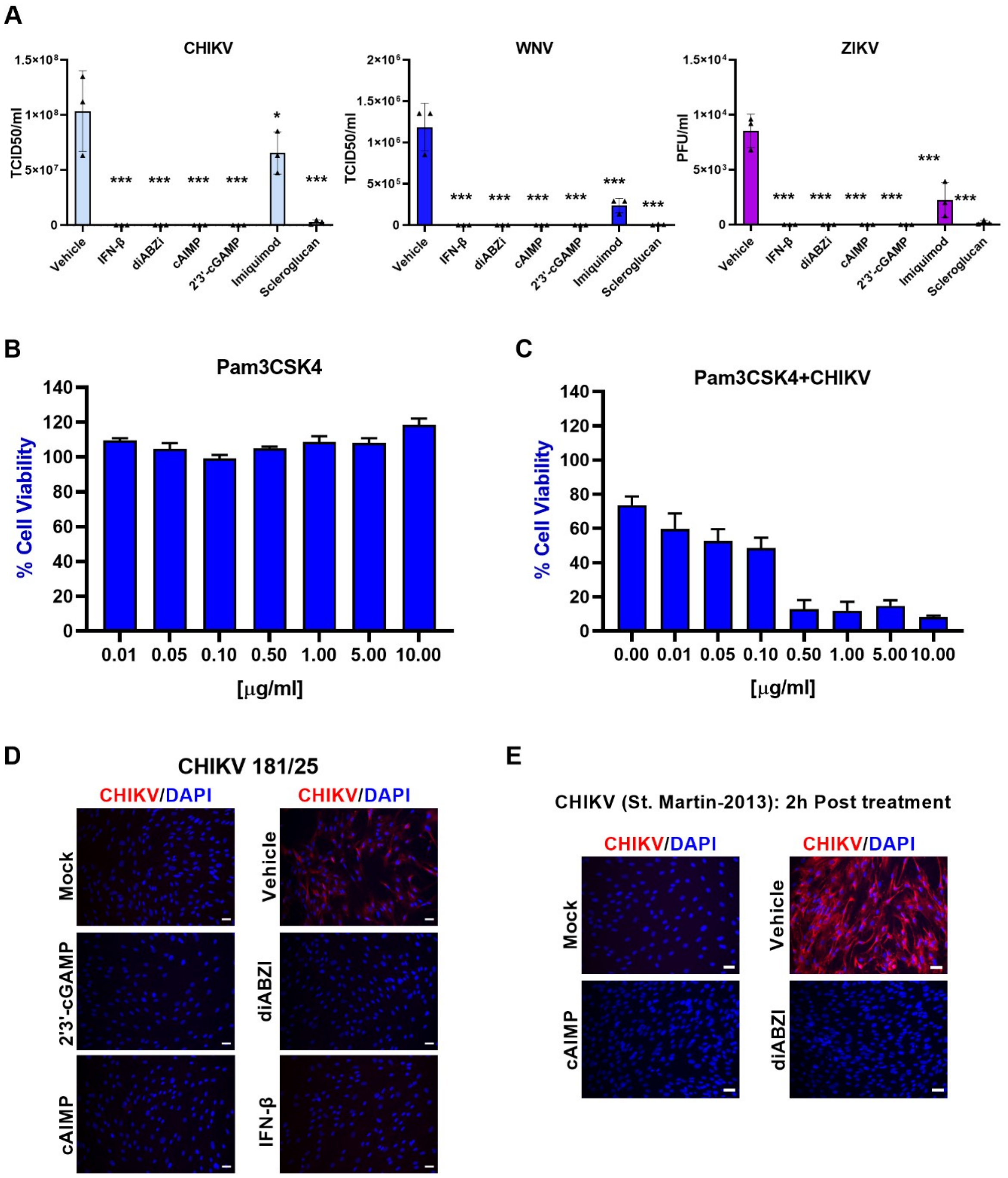

#### Supplementary Figure 3

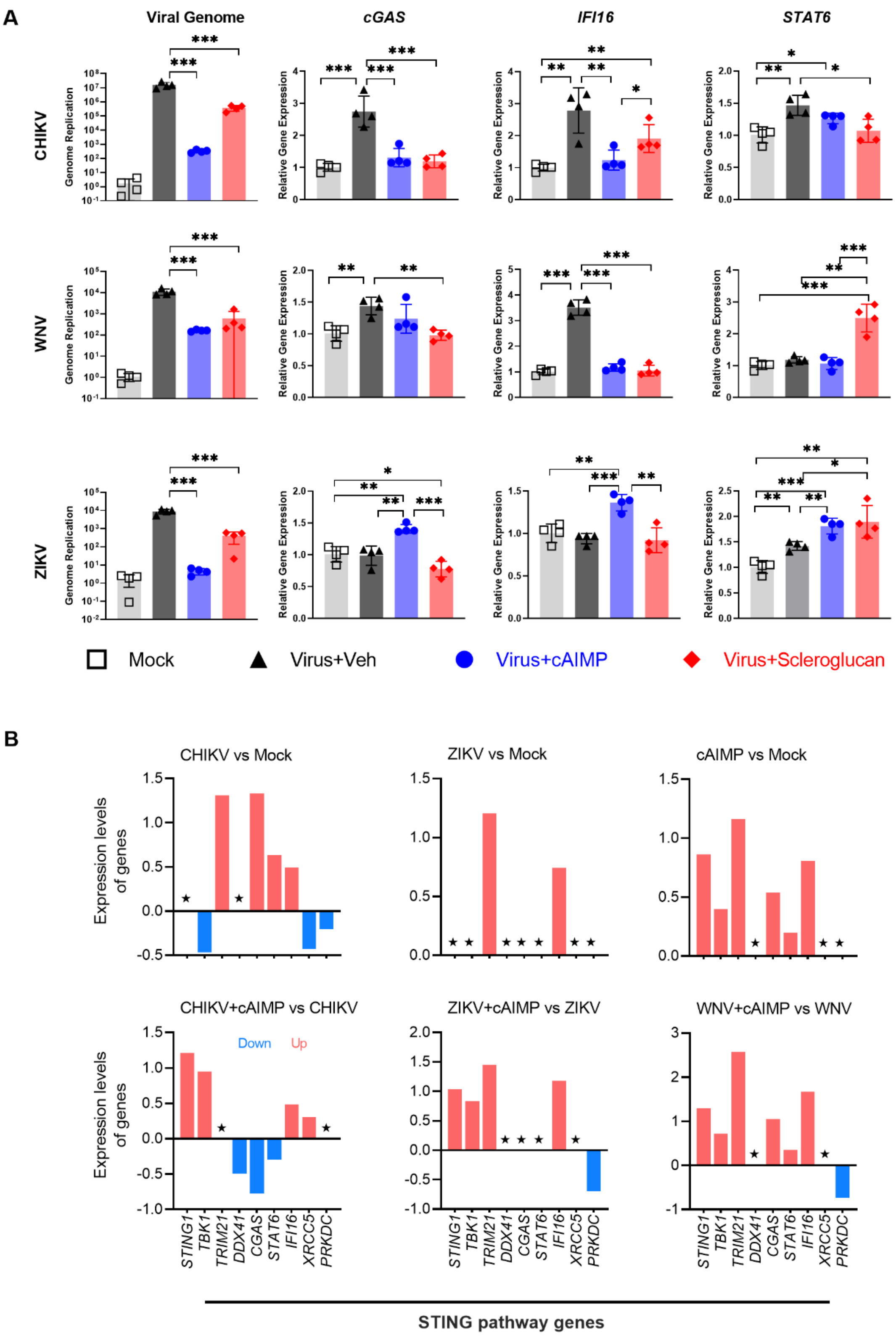

### Supplementary Figure 4

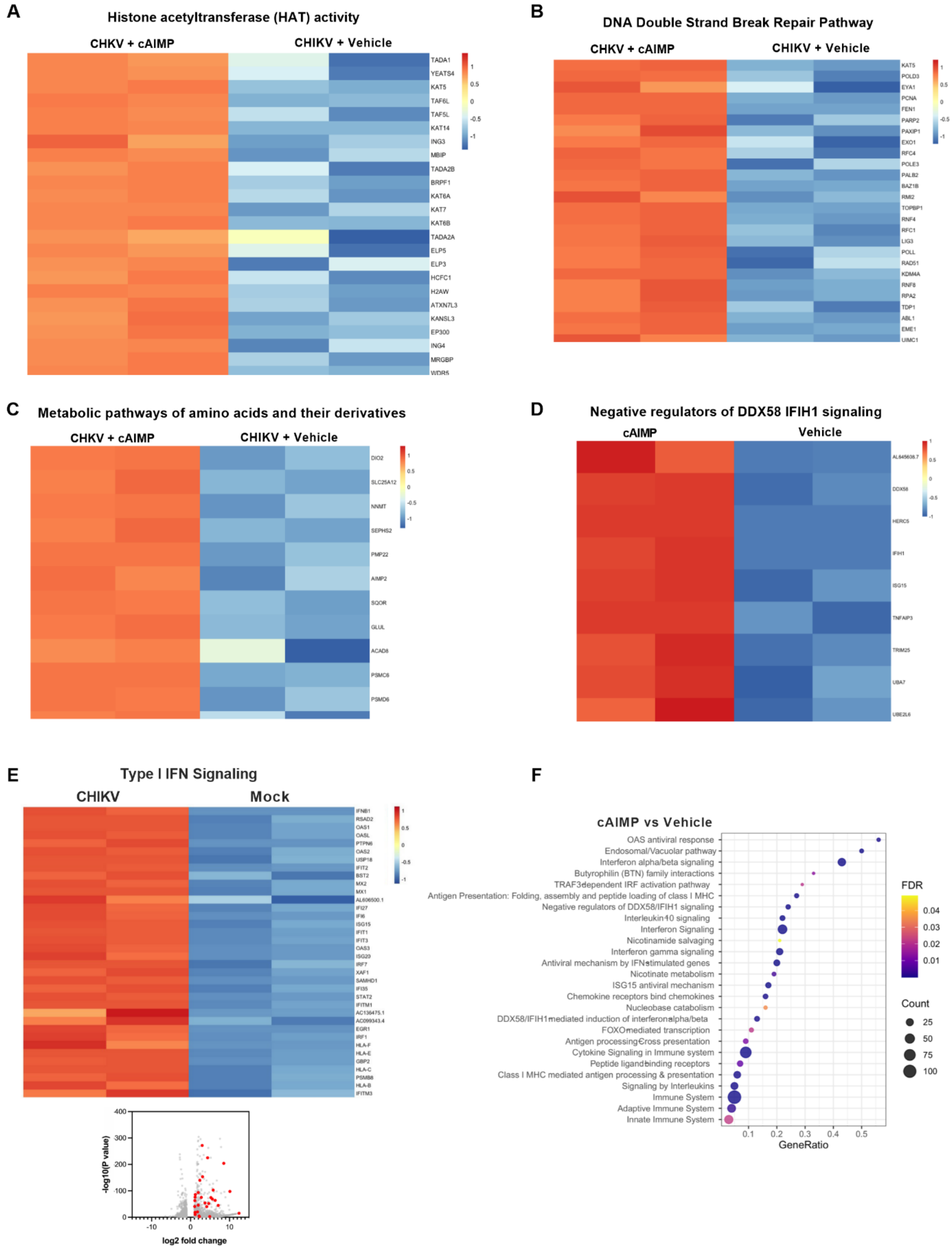

**Supplementary Figure 5**

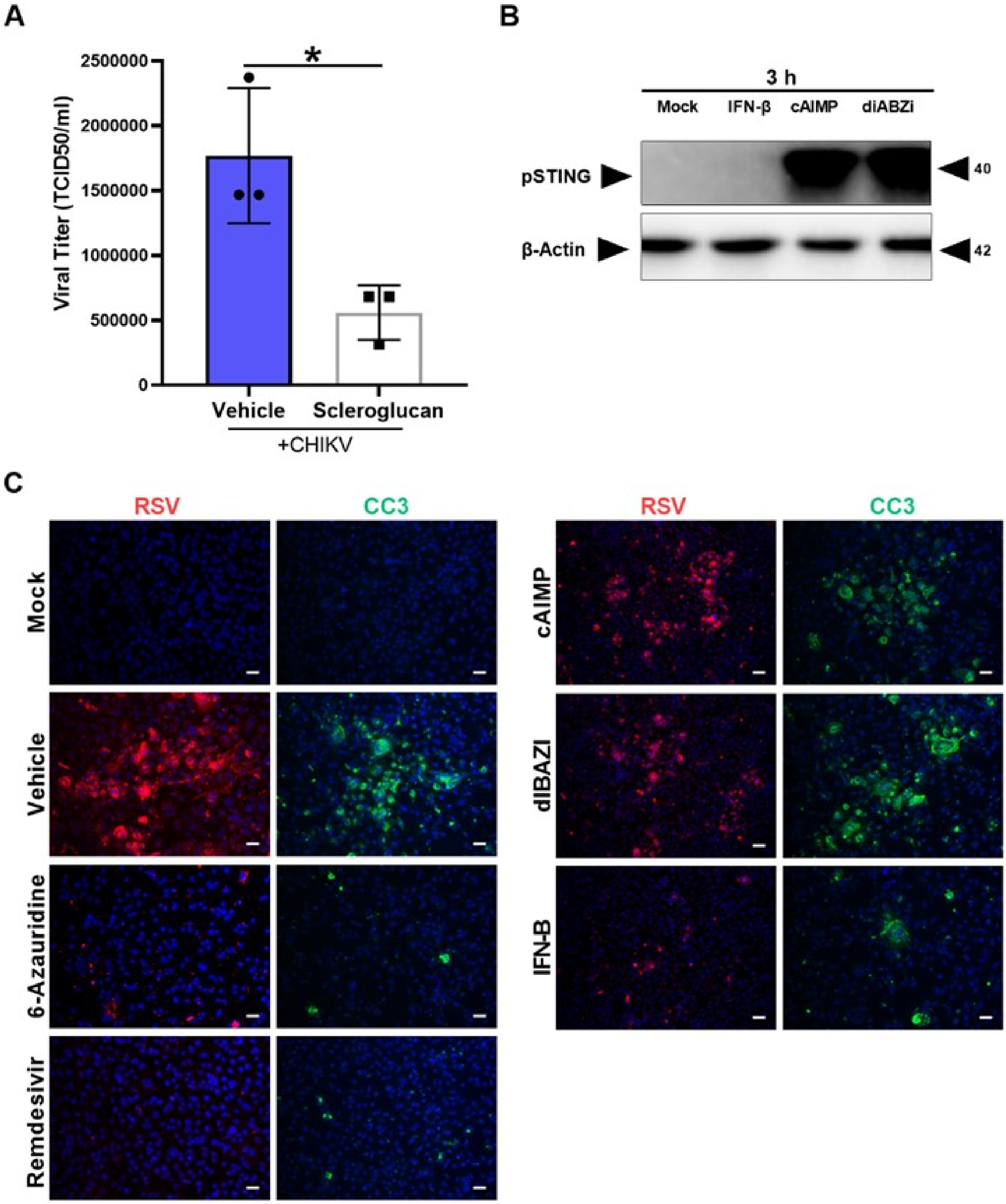


.

### Supplementary Figure 6

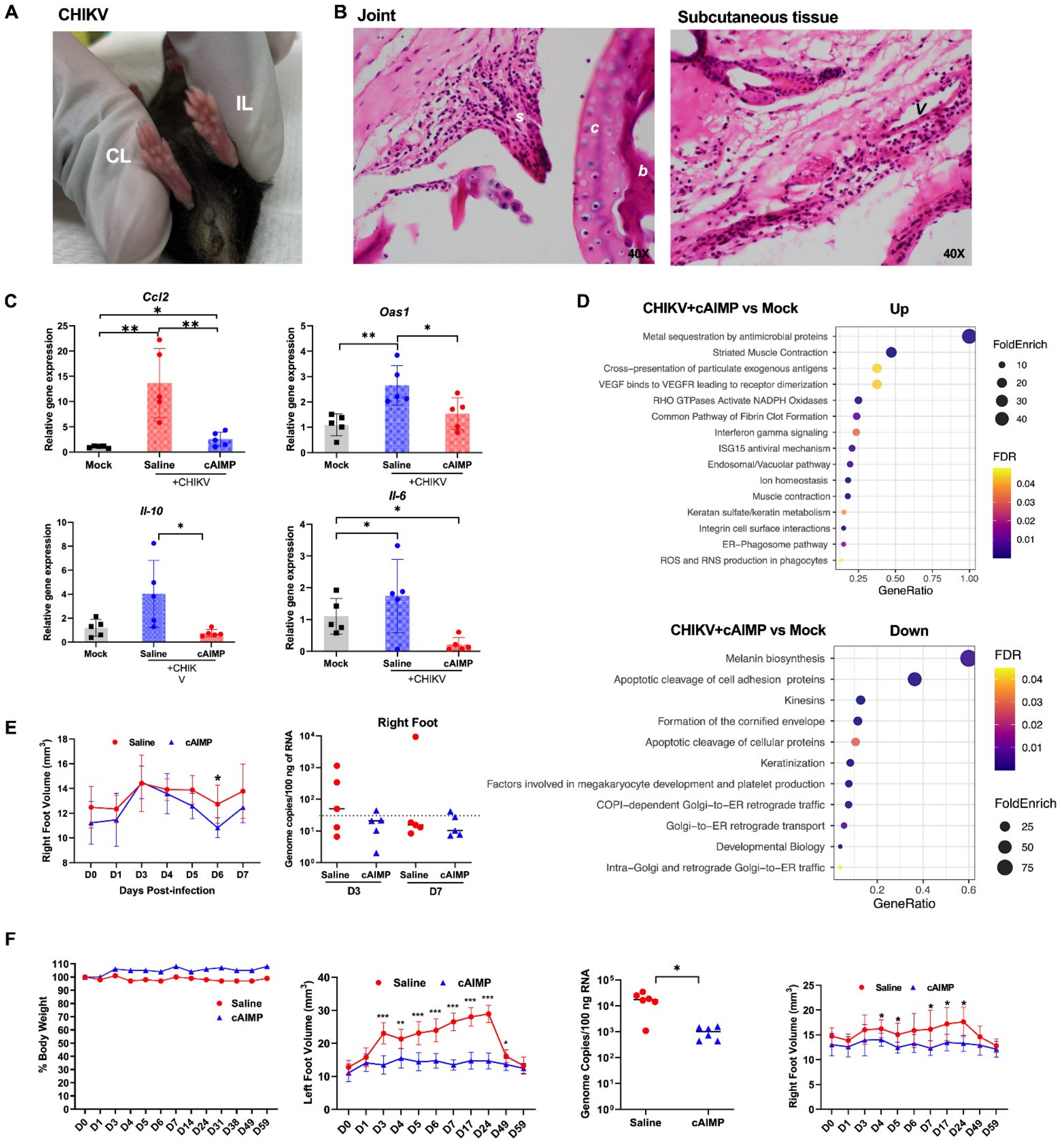

1 **Supplementary Figure 7**

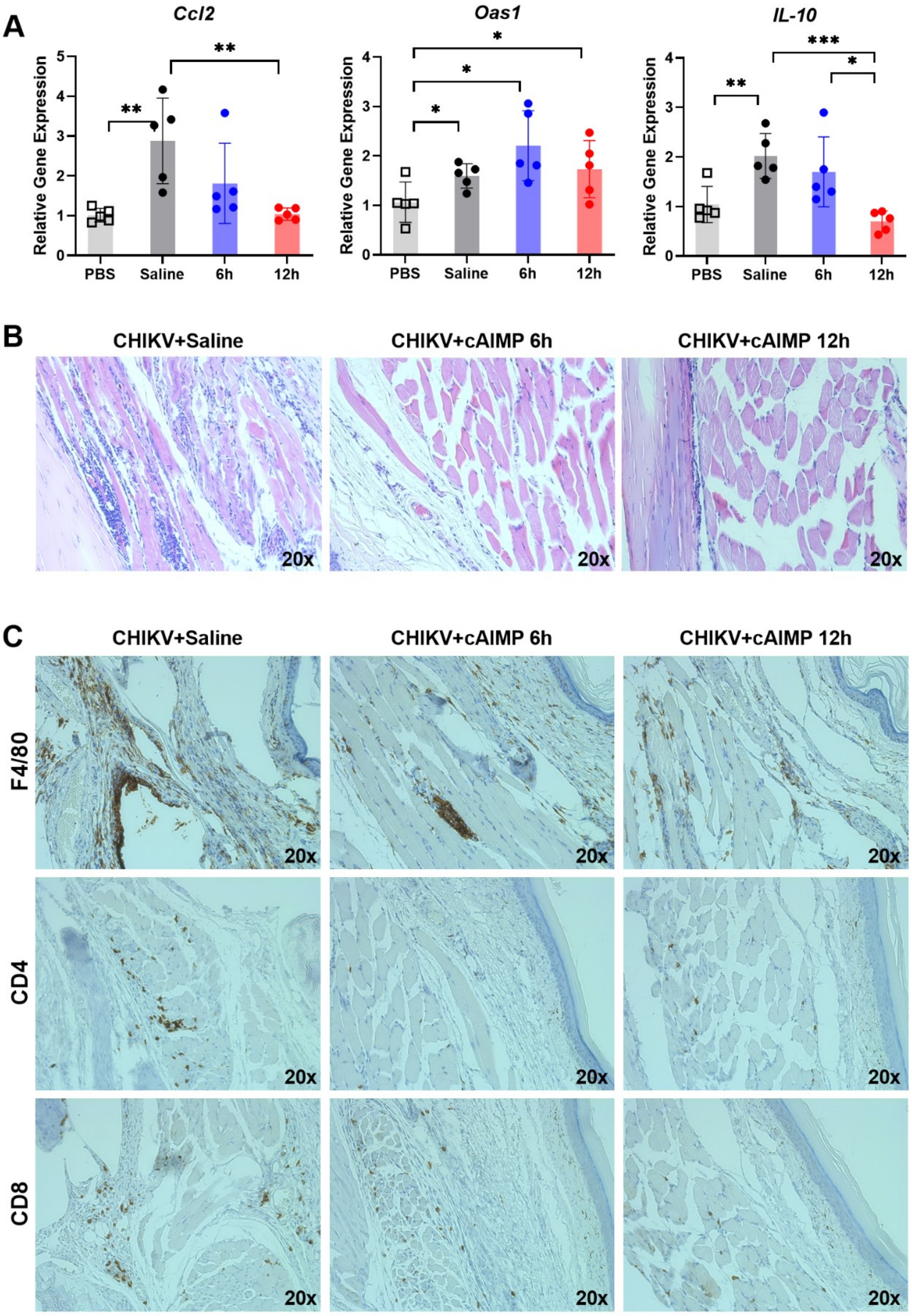

**Supplementary Figure 8**

A

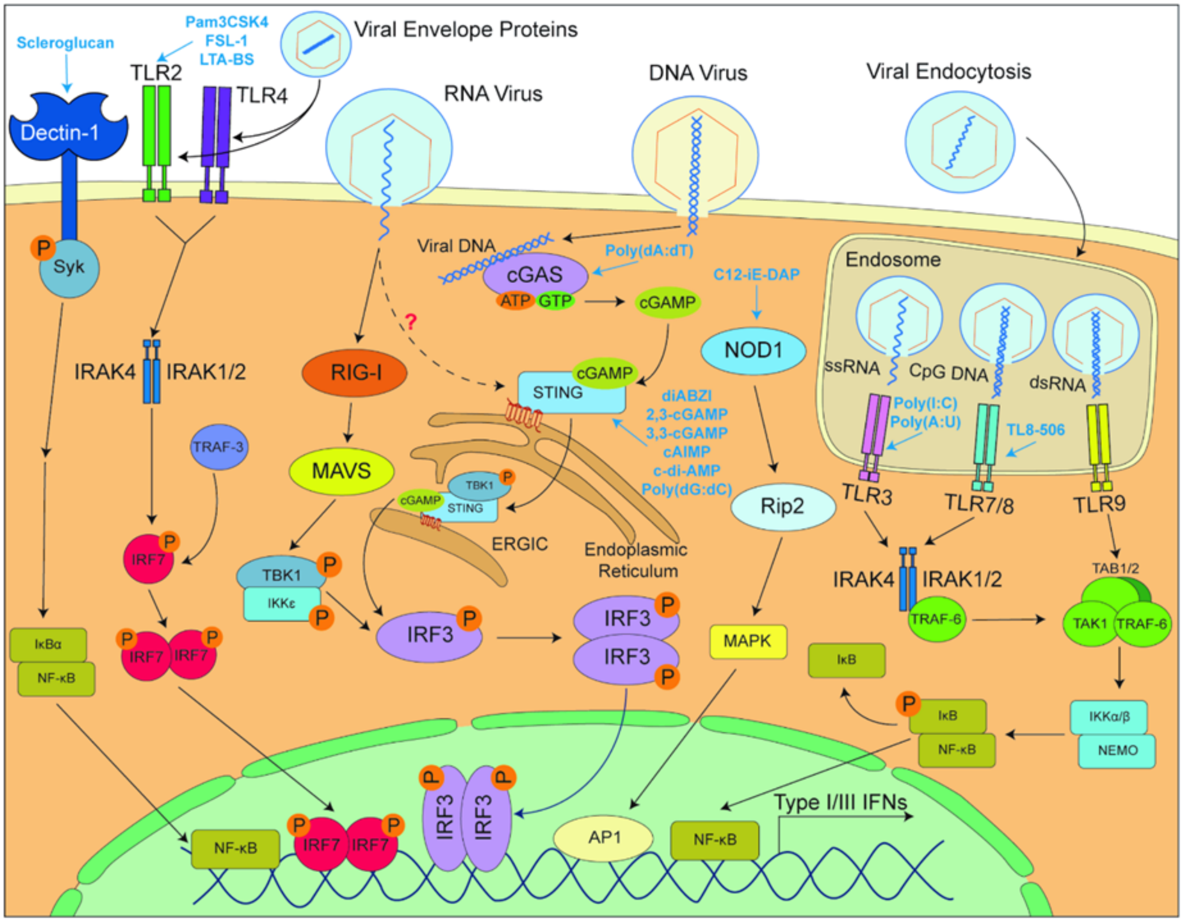
